# M-CSF–stimulated alveolar macrophages safeguard from invasive aspergillosis

**DOI:** 10.64898/2026.07.06.736478

**Authors:** Dalia Sheta, Zeinab Mokhtari, Marlene Strobel, Yidong Yu, Pia Wittmann, Zahraa Abboud, Michael A. G. Kern, Jorge Amich, Nora Trinks, Sebastian Reinhard, Sina Hirsch, Ivan Aleksic, Vasileios Drosos, Eslam S. Ibrahim, Kerstin Günther, Knut Ohlsen, Martin Fraunholz, Christian Stigloher, Albert Garcia Lopez, Sascha Schäuble, Natalie Nieuwenhuizen, Tobias Köhler, Oliver Kurzai, Emmanuel Antoine Saliba, Panagiota Arampatzi, Alexander J Westermann, Paul M. Jordan, Oliver Werz, Jürgen Löffler, Gianni Panagiotou, Hermann Einsele, Markus Sauer, Katrin G. Heinze, Manfred B. Lutz, Heike M. Hermanns, Ulrich Terpitz, Andreas Beilhack

## Abstract

Invasive pulmonary aspergillosis (IPA) is a life-threatening complication in immunocompromised individuals, including recipients of allogeneic hematopoietic cell transplantation (allo-HCT). While systemic neutropenia is traditionally considered the primary risk factor for IPA, we demonstrate that tissue-resident alveolar macrophages (AMs), rather than recruited neutrophils, dictate survival during the critical early window after transplantation. Utilizing an ultra-low dose *Aspergillus fumigatus* infection model that mimics physiological exposure, we identify alveolar macrophages (AMs) as key players in pulmonary antifungal defense. In immunocompromised mice, AMs conferred protection against lethal invasive aspergillosis by day 6, but not day 4 post-allo-HCT. To enhance AM function at the earlier time point, we tested cytokine-based interventions and show that M-CSF, but not IL-34, which both bind to the CSF-1 receptor, promotes migratory activity, phagolysosomal function and fungal killing in both mouse and human primary tissue-resident AMs. In allo-HCT recipient mice, M-CSF treatment preserved lung tissue integrity, suppressed pro-inflammatory cytokines, and protected mice from lethal invasive aspergillosis. The M-CSF-driven protective effect was abrogated upon AM depletion. Our findings demonstrate a critical role of tissue-resident AMs in pulmonary antifungal immunity and suggest that therapeutic modulation of AM activity via M-CSF may offer a promising strategy to combat severe fungal infections in immunocompromised patients.

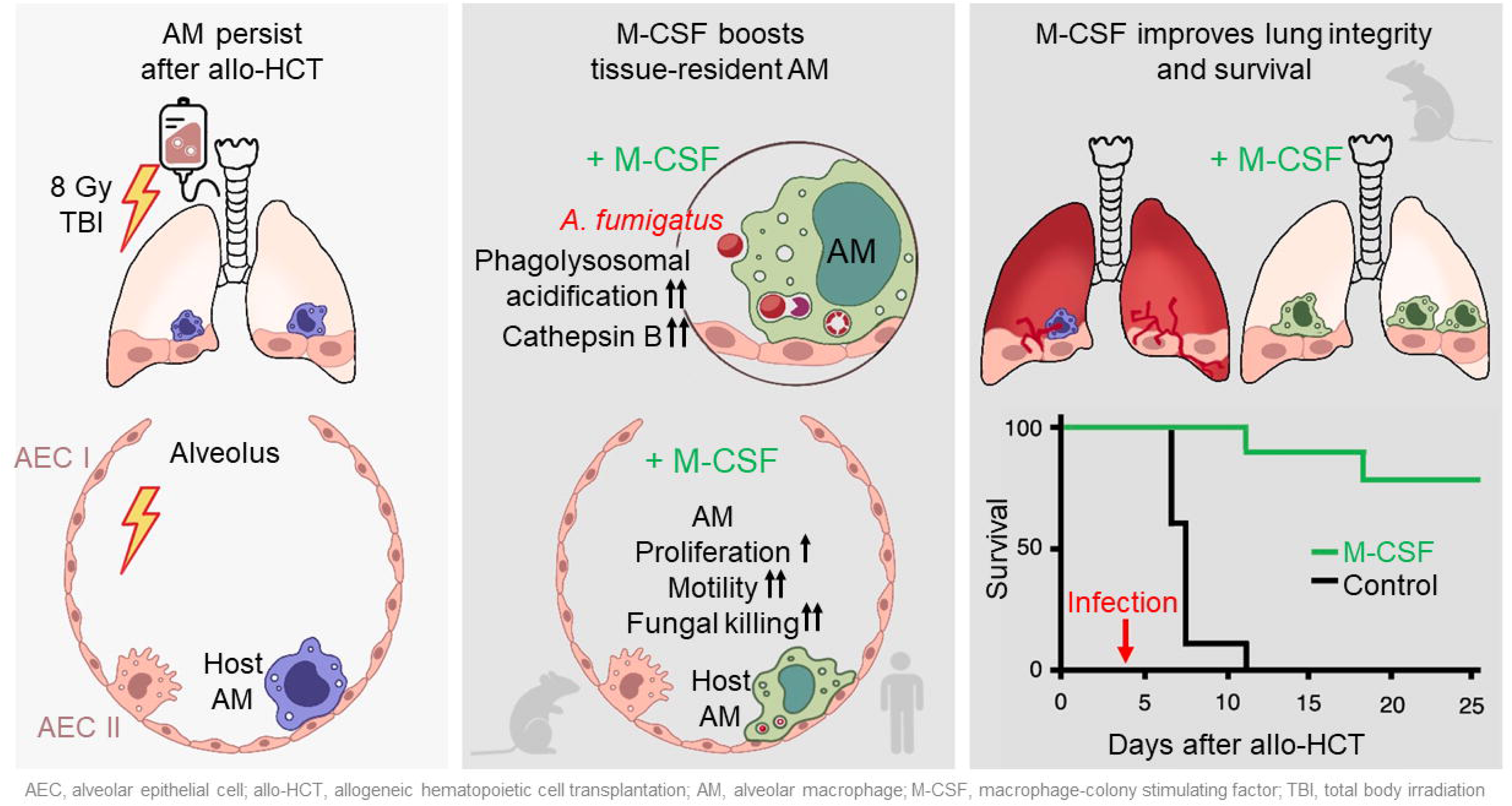

## Introduction

Airborne pathogens pose a persistent clinical threat, particularly to immunocompromised individuals. One of these pathogens is *Aspergillus fumigatus*, a ubiquitous environmental mold and the leading cause of invasive pulmonary aspergillosis (IPA)^1–3^. IPA is a life-threatening infection that primarily affects patients with impaired immunity, such as recipients of allogeneic hematopoietic cell transplantation (allo-HCT)^4–7^. Reported incidence of IPA in allo-HCT recipients ranges from 5% to 15%, depending on study cohort, antifungal prophylaxis, and underlying risk factors^8^. The small size of *A. fumigatus* spores (2–3 μm) enables them to bypass upper airway defenses and deposit deep in the alveoli, where they can germinate and invade lung tissue, leading to a spectrum of diseases from allergic reactions to fulminant IPA. In immunocompetent hosts, alveolar defenses—including epithelial barriers and innate immune cells—rapidly neutralize these spores^9–11^. However, allo-HCT patients are particularly susceptible to IPA due to factors such as neutropenia, delayed immune reconstitution, conditioning regimens, and graft-versus-host disease (GvHD)^12,13^. Notably, IPA incidence follows a bimodal pattern: early cases occur during neutropenia, while later cases are linked to GvHD-associated immunosuppression^14–17^. While systemic immunity can be partially restored using granulocyte colony-stimulating factor (G-CSF)^18–20^, the kinetics and functionality of pulmonary immune recovery post-HCT remain poorly defined^21,22^. This is attributable in part to the limitations of current in vitro systems, which often fail to recapitulate the complexity of the lung microenvironment during post-transplant recovery.

The initial defense against inhaled pathogens^23^, prior to the recruitment of polymorphonuclear neutrophils (PMNs)^24–26^, is provided by the pulmonary epithelium and tissue-resident alveolar macrophages (AMs)^27,28^. The epithelium acts as both a physical barrier and an immune modulator by expelling microorganisms and releasing cytokines, chemokines, and antimicrobial factors that shape downstream responses^29–31^. Yet, *A. fumigatus* spores often bypass these defenses. They invade the alveoli, transform into invasive fungal hyphae, which are capable of breaching tissues and disseminating hematogenously ^30,31^. Under steady state conditions, tissue-resident AMs maintain pulmonary homeostasis through immune surveillance, regulation of inflammation and participation in tissue remodeling and repair in both mice and humans^32,33^. However, the dynamics of tissue-resident immunity in post-transplant lungs remain largely uncharacterized. This gap is particularly striking for AMs, because they are long-lived, self-renewing sentinels strategically positioned to patrol alveolar surfaces, clear inhaled pathogens, and orchestrate local immune responses^27,34,35^. Given their unique location at the pathogen–host interface, AMs represent an attractive but unexplored target for therapeutic modulation after allo-HCT. Whether and how their antifungal capacity can be pharmacologically enhanced in vulnerable patients is therefore a pressing and clinically relevant question.

To address this question, we turned our attention to colony-stimulating factors (CSFs), which have emerged as powerful regulators of myeloid cell biology. Research has focused predominantly on granulocyte-macrophage colony-stimulating factor (GM-CSF) and G-CSF, both of which drive systemic myeloid recovery, neutrophil recruitment, and differentiation^36–38^. By contrast, macrophage colony-stimulating factor (M-CSF) is best known as a critical driver of monocyte survival, differentiation and proliferation^39,40^. However, its role in shaping the function of lung-resident AMs has remained unexplored. This is a crucial gap, as AM biology is not interchangeable with that of circulating monocytes or other tissue-resident macrophages: their longevity, epithelial crosstalk, and unique metabolic and transcriptional programs suggest that they may respond to cytokine signals in fundamentally distinct ways.

To better understand pulmonary immunity against fungal infections, we employed an allo-HCT mouse model combined with intratracheal *A. fumigatus* inoculation. Unlike prior studies relying on supraphysiologic fungal doses or hyper-virulent strains that provoke transient lung inflammation^41,42^, our approach mimics clinically relevant infection kinetics. Our research reveals a previously underappreciated central role of AMs in controlling fungal invasion. We also show that M-CSF enhances AM function by stimulating local AM proliferation, increasing AMs motility, and boosting phagolysosomal activity, thereby promoting fungal killing in both murine and primary human cells. In vivo, M-CSF preserves lung integrity, reduces pro-inflammatory cytokine levels, and improves survival following fungal challenge. Depletion experiments confirm that these protective effects of M-CSF are AM-dependent, highlighting its potential as a promising therapeutic strategy to strengthen frontline pulmonary defenses in immunocompromised hosts.

## Results

### AMs protect against pulmonary *A. fumigatus* infection at day 6, but not day 4 after allo-HCT

To mimic invasive aspergillosis in immunocompromised patients undergoing allo-HCT and to define the window of vulnerability, we challenged an allo-HCT mouse model (C57BL/6→BALB/c) with intratracheal *A. fumigatus* infection. We administered 5×10^4^ spores of the ATCC46645 strain of *A. fumigatus* either 4 days or 6 days after myeloablative total body irradiation and allo-HCT **(Fig. 1A)**. Host survival strictly depended on the timing of the fungal challenge. Only 20% of mice survived infection on day 4 after allo-HCT, whereas day+6 infection resulted in 100% survival (**Fig. 1B).** Consistent with this survival rates, mice infected on day 4 exhibited a significantly higher pulmonary fungal burden than those infected on day 6 **(Fig. 1C, Fig. S1)**. To identify the specific cell types driving this day+6-protective phenotype, characterized the pulmonary innate immune responses at early (24 h) and late (72 h) post-infection times. Flow cytometry analysis of whole lung homogenates and light-sheet fluorescence microscopy (LSFM)^43^ of ∼300-400 μm^2^ lung tissue revealed that AMs outnumbered PMN and monocytes at both time points, with this difference reaching statistical significance by 72 hours **(Fig. 1D, 1E, Fig. S2)**. AMs also displayed a higher proliferation rate than PMN and monocytes **(Fig. 1F)**. To evaluate functional efficacy, we utilized the fluorescent *Aspergillus* reporter (FLARE) system^44^, to differentiate associated (A633^+^) live (dsRed^+^) conidia from killed conidia (dsRed^-^). FLARE analysis demonstrated that AMs achieved significantly higher fungal killing rates 72 h post-infection compared to both PMN and monocytes **(Fig. 1G, H)**.

**Figure 1:**
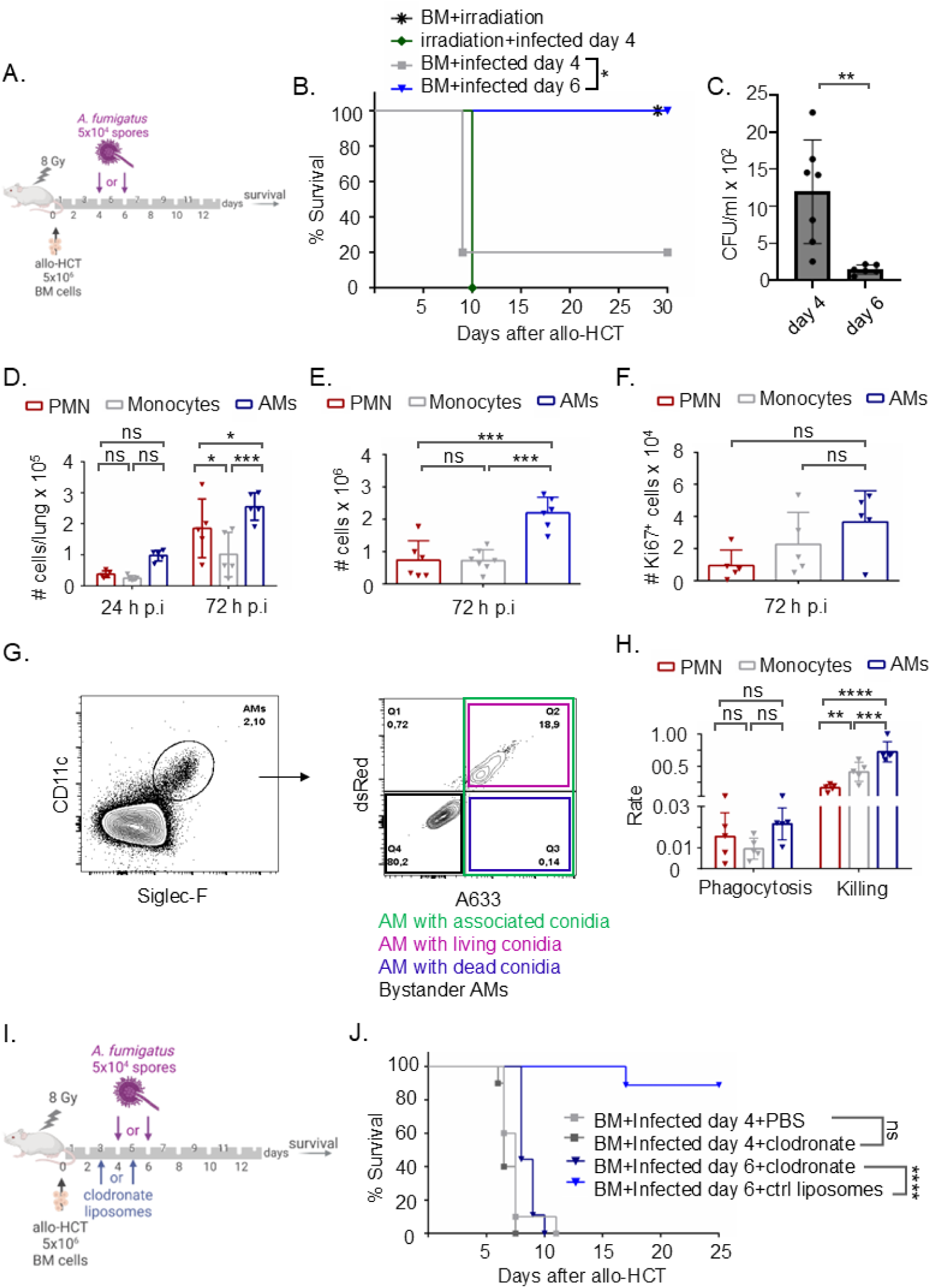
Pulmonary immune response against *A. fumigatus* infection at day 6 after allo-HCT. (**A**) Experimental scheme for *A. fumigatus* intratracheal infection after irradiation with a dose of 8 Gy and allo-HCT of 5×10^6^ BM cells. The infectious dose is 5×10^4^ ATCC46645 *A. fumigatus* spores per mouse at day 4 or day 6 after allo-HCT. (**B**) Survival of mice infected with 5×10^4^ *A. fumigatus* spores at day 4 or at day 6 after irradiation and allo-HCT, in comparison to transplantation, and infection controls. *n* = 10 per group. Statistical significance was determined by Mantel-Cox test (*, *p* ≤ 0.05). (**C**) Quantification of fungal colonies recovered from lung homogenates of mice infected at day 4 or day 6 after allo-HCT. *n* = 6-7. Statistical significance was determined by unpaired T-test. Values were displayed as mean ± SD. (**, *p* ≤ 0.01). (**D**) Flow cytometry quantification of AMs, monocytes and neutrophils numbers at 24 h and 72 h after *A. fumigatus* infection. Statistical significance was determined by two-way ANOVA with Holm-Sidak’s multiple comparisons test. *n* = 5 per group. (**E**) LSFM quantification of AMs, monocytes and neutrophils numbers at 72 h after *A. fumigatus* infection. We performed quantification on 3 different z-stacks, each with a size of 250-400 µm^3^ per mouse, in total 3 mice per condition. *n* = 3. Statistical significance was determined by ordinary one-way ANOVA with Holm-Sidak’s multiple comparisons test. (**F**) Flow cytometry quantification of AMs, monocytes and neutrophils proliferation via Ki67^+^ intracellular staining at 72 h after *A. fumigatus* infection. *n* = 5 per group. Statistical significance was determined by ordinary one-way ANOVA with Holm-Sidak’s multiple comparisons test. (**G**) Contour plot for flow cytometry identification of AMs, which we identified by gating on living cells then single cells followed by SiglecF and CD11c expression. Representative contour plot for phagocytosis and killing of FLARE conidia by AM based on intrinsic expression of dsRed viability signal and surface expression of Alexa Flour 633 (AF633). (**H**) Flow cytometry quantification of phagocytosis and killing rate of FLARE conidia at 72 h after infection. We calculated the rate by dividing the absolute number of cells with engulfed living or dead conidia per the total number of cells of the same type to give phagocytosis and killing rate, respectively. *n* = 5 per group. Statistical significance was determined by two-way ANOVA with Holm-Sidak’s multiple comparisons test. (**I**) Experimental scheme for AM depletion and intratracheal *A. fumigatus* infection with a dose of 5×10^4^ ATCC46645 spores per mouse either at day 4 or at day 6 after irradiation with a dose of 8 Gy and allo-HCT of 5×10^6^ BM cells. We depleted AMs by intratracheal administration of 50 μl of 5 mg/ml clodronate liposomes 24 h before infection. (**J**) Survival of AM-depleted mice infected with *A. fumigatus* spores either at day 4 or at day 6 after irradiation and allo-HCT, in comparison to control liposomes-treated mice and infection controls. *n* = 9-11. Statistical significance was determined by Mantel-Cox test. Values were displayed as mean ± SD. (ns, not significant; *, *p* ≤ 0.05; **, *p* ≤ 0.01; ***, *p* ≤ 0.001, ****, *p* ≤ 0.0001).

Notably, while the absolute numbers of AMs, PMNs, or monocytes did not differ significantly between day 4 and day 6 post allo-HCT **(Fig. S3A, D, F)**, the fungal killing capacity of AMs was profoundly impaired on day 4 **(Fig. S3B, C)**. PMN and monocytes showed no detectable phagocytic activity at this earlier time point **(Fig. S3E, G)**. Together, these data demonstrate that AMs are the primary effector cells defending against *A. fumigatus* after allo-HCT, but their functional capacity is temporarily suppressed during the immediate post-transplantation window.

### Local depletion of alveolar macrophages abrogates post-transplant protection

Having established that AMs correlate with protection on day 6, we next sought to determine we next sought to determine whether this local tissue-resident pool is strictly required for host survival. We selectively depleted Ams via a single intratracheal administration of clodronate liposomes (50 µl of 5 mg/ml). This approach effectively eliminated AMs within 24 hours while sparing other innate immune populations within the bronchoalveolar lavage (BAL) fluid **(Fig. S3H)**. Following *A. fumigatus* challenge on day 6 post-allo-HCT, AM-depleted mice completely failed to survive the infection, in contrast to non-depleted controls **(Fig. 1I, J)**. Crucially, this loss of protection occurred despite the normal, unhindered recruitment and recovery of alternative innate effectors, such as neutrophils and monocytes, in the lungs **(Fig. 1D, E)**. These loss-of-function experiments confirm that tissue-resident AMs are indispensable for pulmonary defense against *A. fumigatus* following allo-HCT. Furthermore, because this protective capacity emerges only after a spontaneous 6-day recovery period, these findings suggest that exogenous cytokine stimulation might be leveraged to accelerate AM maturation and restore frontline defense during the early window of vulnerability.

### M-CSF enhances AM motility in vitro

While activation and M1/M2 polarization pathways of monocyte-derived macrophages are well characterized^45,46^, the phenotypic modulation of embryonic-derived, niche-dependent, tissue-resident AMs by cytokines remains poorly understood. Among niche-specific signals, macrophage colony-stimulating factor receptor (CSF1R) and its ligands—M-CSF and interleukin-34 (IL-34)—are critical^47,48^, but their specific impact on AM functional phenotypes have not been delineated^49^.

We initially explored how M-CSF and IL-34 influence AM motility, a requirement for alveolar immune surveillance. To mimic the alveolar niche, we established a tri-culture system with primary alveolar epithelial cells (AECs), naïve AMs, and fluorescent tdTomato^+^-labelled *A. fumigatus* conidia. AECs delayed conidial germination (**Fig. S4A, B**), an effect marginally extended by AMs (**Fig. S4C**). Notably, AECs enhanced AM motility; AMs in the co-culture exhibited longer migration paths **(Fig. S4D)**, higher diffusion coefficients **(Fig. S4E)** and increased speed **(Fig. S4F)** compared to AM single cultures with fungal spores. In presence of AECs, AMs also exhibited broader turning angles **(Fig. S4G)**, indicating random, non-directed movement. Consequently, we utilized this physiological co-culture platform for subsequent cytokine assays. Next, we pre-treated AMs in vitro with M-CSF or IL-34. Despite sharing the CSF1R receptor, only M-CSF enhanced AM motility **(Fig. 2)**. Compared to PBS **(Video S1)** or IL34 **(Video S3)**, M-CSF-stimulated AMs migrated longer distances **(Fig. 2A, Video S2),** exhibited higher diffusion coefficients **(Fig. 2B)**, moved faster **(Fig. 2C)** and showed broader turning angles **(Fig. 2D)**. M-CSF consistently outperformed other cytokines, including GM-CSF, G-CSF, and IL-3, M-CSF **(Fig. S5A-D)**. This response was not observed in other tissue-resident macrophages, like peritoneal macrophages **(Fig. S5E-H)**, underscoring an AM-specific sensitivity to M-CSF.

**Figure 2:**
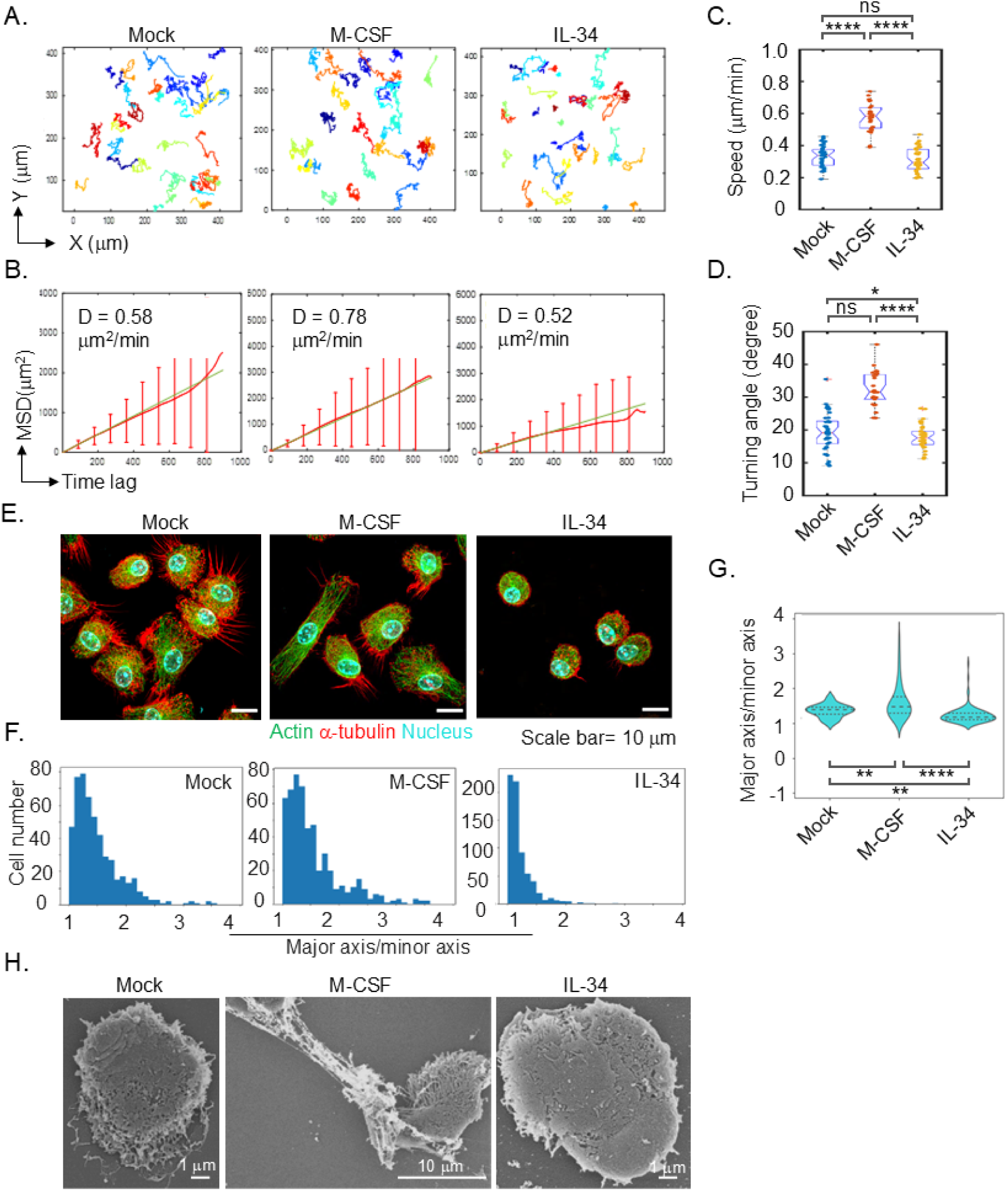
Differential effects of M-CSF and IL-34 on AMs upon in vitro stimulation. (**A**) Display of the migration path of cytokine pre-stimulated AMs while co-cultured with *A. fumigatus* ATCC46645 spores and underlying AEC. Imaging was performed with confocal microscope for 15 h. AMs were stimulated with 300 U/mL of M-CSF and 6×10^3^ U/mL of IL-34. Number of cells used for analysis is 40-50 cells. (**B**) Quantification of MSD and diffusion coefficient of AMs based on AM migration in (A). Number of cells used for analysis is 40-50 cells. (**C, D**) Quantification of the migration speed (C) and turning angle distribution (D) of AMs from (A). Number of cells used for analysis is 40-50 cells. Statistical significance was determined by ordinary one-way ANOVA with Holm-Sidak’s multiple comparisons test. (**E**) Representative SIM images of AMs after in vitro M-CSF and IL-34 stimulation for 48 h. We visualized cell cytoskeleton using phalloidin Atto643, primary mouse anti α-tubulin to which a secondary goat anti mouse F(ab)2-fragment.A488 was used and stained the nucleus with Hoechst 34580. Scale bars = 10 µm. (**F**) Cell shape distribution from (E) (**G**) Quantification of AM eccentricity after in vitro M-CSF and IL-34 stimulation for 48 h. The quantification is based on 10 images per condition (25-30 cells per image) for one experiment with a total of 3 different experiments. *n* = 3. Statistical significance was determined by ordinary one-way ANOVA with Holm-Sidak’s multiple comparisons test. (**H**) Representative SEM images for AMs after in vitro M-CSF and IL-34 stimulation for 48 h. Values were displayed as mean ± SD. (ns, not significant; *, *p* ≤ 0.05; **, *p* ≤ 0.001, ****, *p* ≤ 0.0001).

To identify the structural basis of this phenotype, we examined AM cytoskeletal organization using structured illumination microscopy (SIM). M-CSF-treated AMs displayed marked elongation of phalloidin-labeled actin filaments and α-tubulin-labeled microtubules (**Fig. 2E**), contrasting sharply with the compact morphology of IL-34-treated cells. Morphometric quantification (**Fig. 2F, G**) and SEM imaging (**Fig. 2H**) confirmed that M-CSF induced a significantly more elliptical cellular shape. Collectively, these findings reveal that M-CSF drives cytoskeletal remodeling in Ams to promote a highly motile, exploratory phenotype, potentially optimizing their ability to clear alveolar pathogens.

### M-CSF boosts lysosomal function and fungicidal activity in both murine and human AMs

Given the crucial role of phagocytosis in antifungal host defense^50^, we evaluated the effects of M-CSF and IL-34 on AM engulfment and microbial killing. Although M-CSF did alter the rate of fungal phagocytosis compared to PBS or IL-34 treatment **(Fig. 3A)**, as confirmed by similar AM-fungal contact under SEM **(Fig. S5I**), it significantly enhanced fungal killing **(Fig. 3B)**. Mechanistically, M-CSF markedly increased acidified lysosomal structures and enhanced cathepsin B proteolytic activity compared to PBS- or IL-34 (**Fig. 3C**). This was evidenced by elevated total cellular fluorescence of Acridine Orange (indicating increased lysosomal acidification required for phagolysosomal maturation) and Magic Red (reflecting enhanced cathepsin B-mediated proteolysis) **(Fig. 3D, E)**. In contrast, reactive oxygen species (ROS) production remained unchanged across treatments **(Fig. 3F, G)**, indicating that M-CSF acts independently of ROS-mediated pathways.

**Figure 3:**
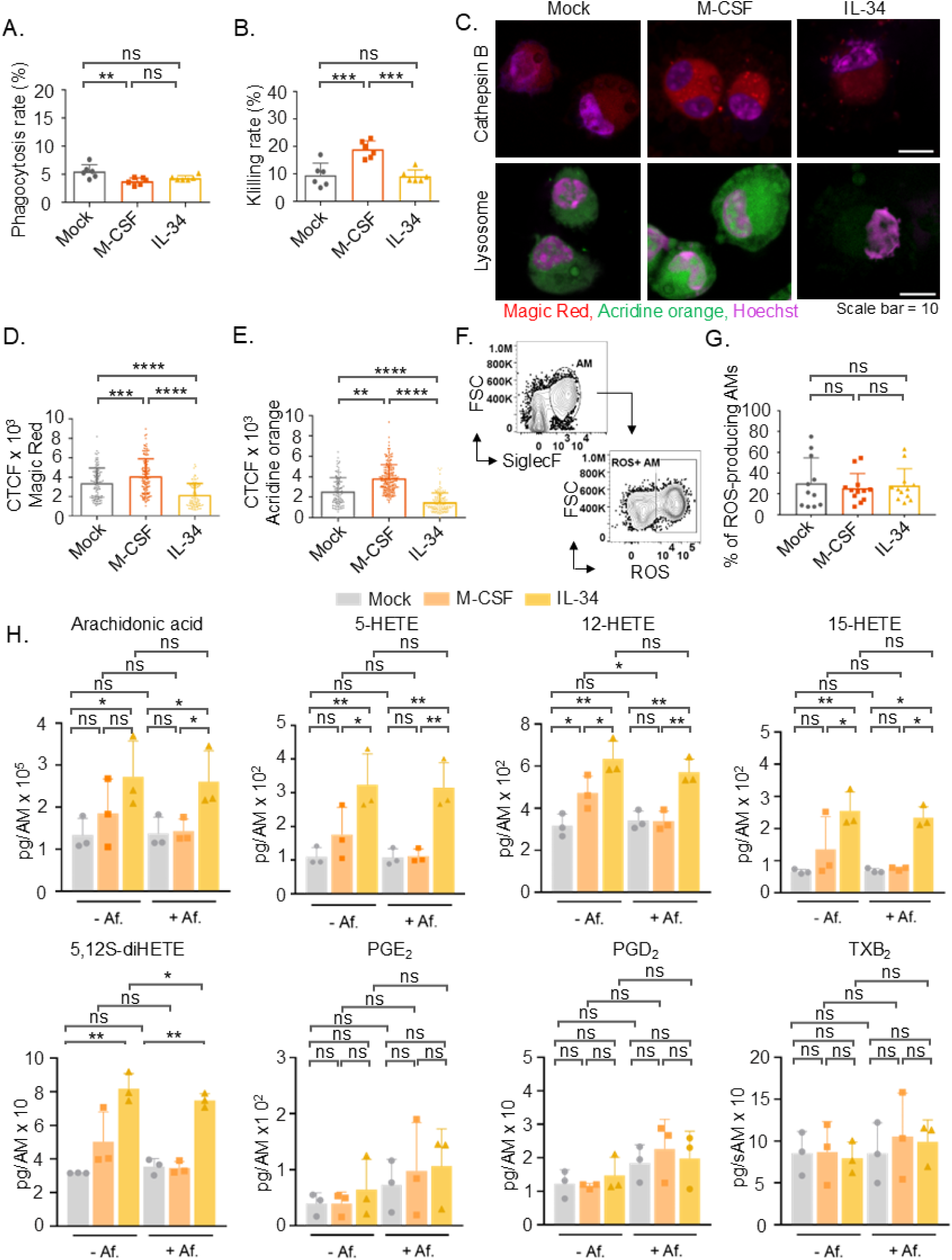
Quantification of AM phagocytosis, killing capacity, and lipid mediator production upon M-CSF and IL-34 in vitro stimulation. (**A, B**) Flow cytometry analysis of phagocytosis rate (A) and killing rate (B) of (1-1.5×10^5^) AMs co-cultured with FLARE conidia and AEC for 1 h in vitro with MOI of 1:1. AMs were isolated by BAL and stimulated with cytokines for 48 h prior to co-culture with conidia. n = 3. Statistical significance was determined by ordinary one-way ANOVA with Holm-Sidak’s multiple comparisons test. (**C**) Confocal fluorescence imaging of the lysosomal acidity marker Acridine Orange and lysosomal cathepsin B enzyme activity using Magic Red on AMs. AMs were stimulated with cytokines for 48 h prior to co-culture for 1 h with ATCC46645 conidia. (**D, E**) Quantification showing corrected total cell fluorescence of (D) cathepsin B-positive cells and (E) acidified lysosomal structures. Total cells for cathepsin B staining n=L106 (Mock), n=L127 (M-CSF-AMs), n= 95 (IL-34-AMs) and for Acridine Orange staining n=L117 (Mock), n=L185 (M-CSF-AMs), n= 145 (IL-34-AMs). Values were displayed as mean ± SD. Statistical significance was determined by ordinary one-way ANOVA with Holm-Sidak’s multiple comparisons test. (**F**) Representative contour plots of flow cytometry analysis of ROS production by AMs. (**G**) Quantification of the percentage of ROS-producing AMs following cytokine pre-stimulation for 48 h and co-culture with *A. fumigatus* ATCC46645 conidia for 6 h. n= 6. Statistical significance was determined by ordinary one-way ANOVA with Holm-Sidak’s multiple comparisons test. (**H**) Lipid mediators analysis of AMs co-cultured with ATCC46645 and pretreated with M-CSF, IL-34 or vehicle in the presence or absence of *A. fumigatus* (+/- Af.) for 6 h. Arachidonic acid release and formed arachidonic acid metabolites, such as 5-,12-, or 15-hydroxyeicosatetraenoic acid (HETE), 5,12-diHETE, prostaglandin (PG) E_2_, PGD_2_ and thromboxane B_2_ were analyzed in the supernatants. *n* = 3. Data, expressed in pg/AM, are given as mean ± SD and log-transformed for statistics. Statistical significance was determined by two-way ANOVA with Tukey’s multiple comparisons test. Values were displayed as mean ± SD. (ns, not significant; *, *p* ≤ 0.05; **, *p* ≤ 0.01; ***, *p* ≤ 0.001, ****, *p* ≤ 0.0001).

Because lipid mediators critically modulate pulmonary inflammation during infection^51,52^, particularly arachidonic acid derivatives which regulate immune cell recruitment and vascular permeability^53,54^, we analyzed their lipid mediator profilew of AM 6 h of co-culture with *A. fumigatus*. Compared to PBS, IL-34–treated AMs displayed elevated levels of arachidonic acid and its pro-inflammatory hydroxylated metabolites, including 5-, 12-, 15-hydroxyeicosatetraenoic acid (HETE) and 5,12S-diHETE regardless of infection status. In contrast, M-CSF did not significantly alter lipid mediator profiles, except for a minor infection-independent increase of 12-HETE and 5,12S-diHETE. Levels of prostaglandin E_2_, prostaglandin D_2_, or thromboxane B_2_ were unaltered across all condition (**Fig. 3H).** Thus, while IL-34 induces pro-inflammatory lipid pathways, M-CSF preferentially enhances lysosomal killing capacity while maintaining a pro-resolving phenotype that limits collateral tissue damage.

To verify that lysosomal acidification drives this enhanced microbial clearance, we measured the survival of an acid-resistant *Staphylococcus aureus* mutant strain^55,56^ within Ams. While the phagocytosis rate of acid-resistant *S. aureus* USA300_JE2 was unaffected by cytokine treatment **(Fig. S5J)**, we recovered significantly more bacterial colony-forming units (CFUs) from M-CSF-treated AM after overnight culture, confirming that M-CSF-mediated protection is subverted by acid-resistant pathogens **(Fig. S5K)**.

To establish the translational relevance of these mechanisms, we primary human AMs isolated from human lung biopsies (huAMs). Replicating our murine data, M-CSF–stimulated huAMs exhibited enhanced motility, showing longer migration paths, higher diffusion coefficients, increased migration speed and broader turning angles compared to PBS- or IL-34 controls (**Fig. 4A–D**). Furthermore, M-CSF increased acidified lysosomal structures and cathepsin B activity in huAMs (**Fig. 4E–G**), culminating in and significantly improved *A. fumigatus* killing despite unaltered phagocytic uptake **(Fig. 4H–J**). Collectively, these findings demonstrate that M-CSF conserves its mechanism across species, augmenting lysosomal proteolytic activity and fungicidal capacity without inducing tissue-damaging ROS or pro-inflammatory lipid cascades.

**Figure 4:**
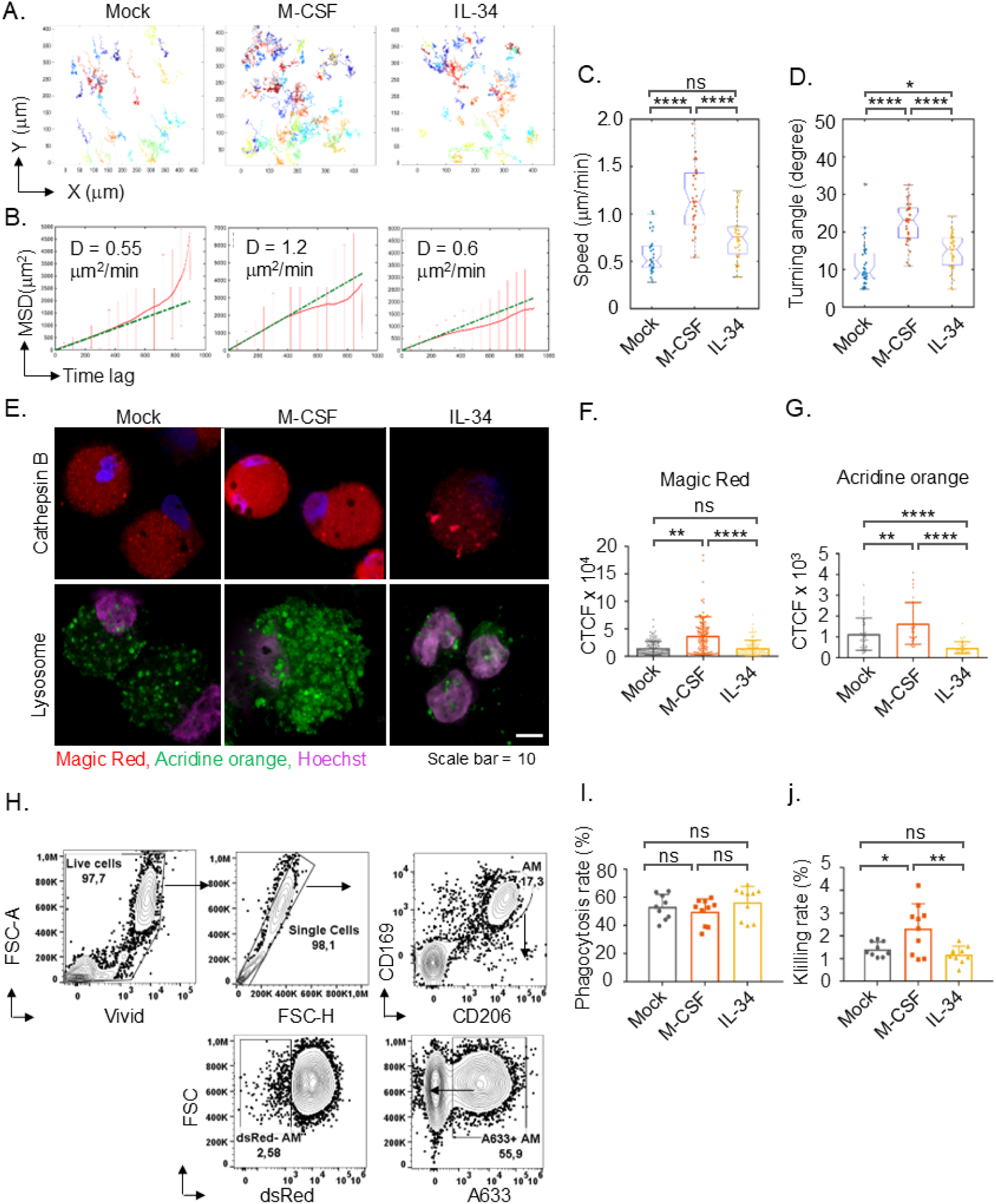
Characterization of huAM migration behavior, phagocytic activity, and killing capacity upon M-SCF and IL-34 in vitro stimulation. (**A**) Display of the migration path of cytokine pre-stimulated huAMs while co-cultured with *A. fumigatus* ATCC46645 spores and underlying AEC. Imaging was performed with confocal microscope for 15 h. huAMs were stimulated with 300 U/mL of M-CSF and 6×10^3^ U/mL of IL-34 for 24 h before imaging. Number of cells used for analysis is 40-50 cells. (**B**) Quantification of MSD and diffusion coefficient of huAMs based on huAM migration in (A). Number of cells used for analysis is 40-50 cells. (**C, D**) Quantification of the migration speed (C) and turning angle distribution (D) of huAMs from (A). Number of cells used for analysis is 40-50 cells. Statistical significance was determined by ordinary one-way ANOVA with Holm-Sidak’s multiple comparisons test. (**E**) Confocal fluorescence imaging of the lysosomal acidity marker, Acridine Orange, and lysosomal cathepsin B enzyme using Magic Red on huAMs. huAMs were stimulated with cytokines for 24 h prior to co-culture for 1 h with ATCC46645 conidia. (**F, G**) Quantification showing corrected total cell fluorescence of (f) cathepsin B-positive cells and (g) acidified lysosomal structures. Total cells for cathepsin B staining n=L164 (Mock), n=L149 (M-CSF-huAMs), n= 111 (IL-34-huAMs) and for Acridine Orange staining n=L53 (Mock), n=L32 (M-CSF-huAMs), n= 45 (IL-34-huAMs). Statistical significance was determined by ordinary one-way ANOVA with Holm-Sidak’s multiple comparisons test. (**H**) Representative contour plots of flow cytometry identification of huAMs isolated from human lung biopsies together with the contour plots for the phagocytosis and killing of FLARE conidia by huAMs based on intrinsic expression of dsRed viability signal and surface expression of A633. (**I, J**) Flow cytometry analysis of phagocytosis rate (i) and killing rate (j) of (4×10^4^) huAMs co-cultured with FLARE conidia (MOI of 1:5) and huAEC for 1 h in vitro. huAMs were isolated from fresh patients’ lung biopsies and stimulated with cytokines for 24 h prior to co-culture with conidia. n = 7. Statistical significance was determined by ordinary one-way ANOVA with Holm-Sidak’s multiple comparisons test. Values were displayed as mean ± SD. (ns, not significant; *, *p* ≤ 0.05; **, *p* ≤ 0.01, ****, *p* ≤ 0.0001).

### Exogeneous M-CSF restores early post-transplant protection and drivel local AM proliferation

Based on our in vitro findings, we reasoned that exogenous M-CSF administration immediately after allo-HCT could fortify the frontline pulmonary response and protect mice during the day 4 window of vulnerability. Using the same allo-HCT model, we intravenously treated mice with M-CSF at –1, 5, and 20 hours relative to allo-HCT, followed by *A. fumigatus* challenge on day 4 **(Fig. 5A)**. To minimize off-target effects of M-CSF, we selected a physiological dose that maintained serum M-CSF levels comparable to healthy controls without altering bone marrow myeloid populations **(Fig S.6A-C).** Furthermore, CSF1R expression in the lung was restricted primarily to AMs and monocytes **(Fig S.6D-F)**; given the scarcity of monocytes during this post-immunosuppression window, this restricted distribution ensures highly localized targeting. Remarkably, M-CSF-treated mice exhibited vastly improved clinical outcomes, achieving 90% survival compared to only 45% in the PBS control group **(Fig. 5B)**.

**Figure 5:**
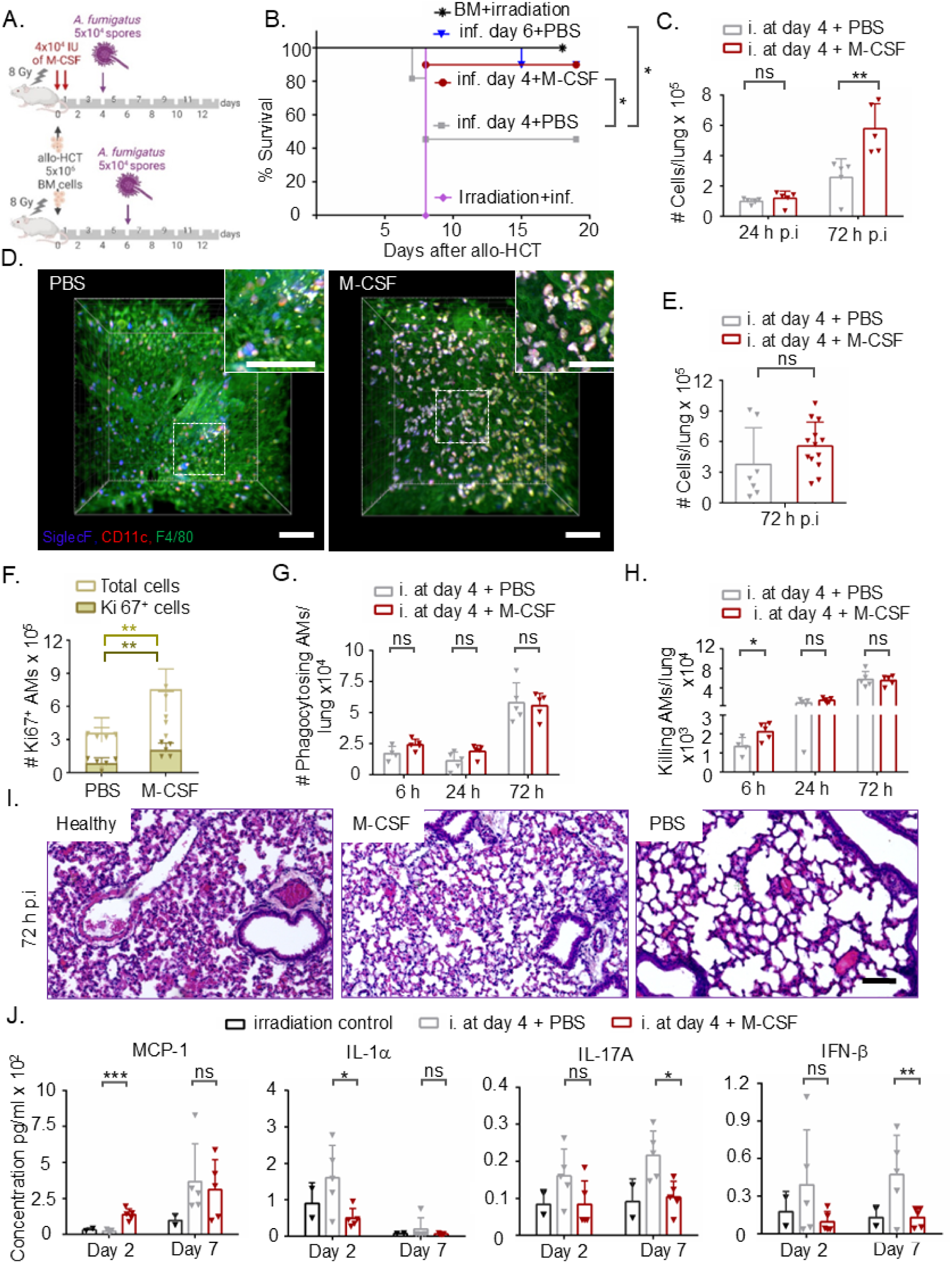
The influence of in vivo M-CSF treatment on AM responses and the outcome of *A. fumigatus* infection. (**A**) Experimental scheme for intratracheal *A. fumigatus* infection with a dose of 5×10^4^ ATCC46645 spores per mouse +/- in vivo M-CSF treatment after irradiation with a dose of 8 Gy and allo-HCT of 5×10^6^ BM cells. We injected M-CSF intravenously in 3 doses with a single dose of 4×10^4^ U/120 µl sterile H_2_O at −1, 5, 20 h of allo-HCT. (**B**) Survival of mice infected with *A. fumigatus* either at day 4 or day 6 after irradiation, allo-HCT and −/+M-CSF treatment in comparison to transplantation and infection controls. Statistical significance was determined by Mantel-Cox test (*, *p* ≤ 0.05). *n* = 5-11 per group. (**C**) Flow cytometry quantification of AM frequency at 24 h and 72 h after *A. fumigatus* infection. *n* = 5 per group. (**D**) LSFM images for AMs interaction with *A. fumigatus* at 72 h after infection. Scale bars = 80 µm. (**E**) LSFM quantification of AM number at 72 h after *A. fumigatus* infection. We quantified 3 different z stacks, each with a size of 250-400 µm^3^, per each mouse with total of 3 mice per condition. (**F**) Flow cytometry analysis of AM proliferation via Ki67^+^ intracellular staining at 72 h after *A. fumigatus* infection at day 4. *n* = 5 per group. (**G, H**) Flow cytometry quantification of absolute numbers of AMs involved in fungal phagocytosis (g) and fungal killing (h) at 6 h, 24 h (day 2) and 72 h (day 7) after *A. fumigatus* infection. *n* = 5 per group. (**I**) Representative H&E-stained lung tissue sections at 72 h after *A. fumigatus* infection. Scale bar = 100 µm, *n* = 3 per group. (**J**) Flow cytometry quantification of serum cytokine and chemokine levels at two different time points before (day 2) and after (day 7) *A. fumigatus* infection. Values were displayed as mean ± SD. Statistical significance was determined by unpaired t-test (*, P ≤ 0.05, **, *p* ≤ 0.01, ***, *p* ≤ 0.001).

To elucidate the mechanisms underlying this protection, we analyzed the local pulmonary innate immune response. At 72 h post-infection, flow cytometry analysis revealed a ∼1.5-fold increase in AM numbers in M-CSF–treated mice compared to controls **(Fig.**LJ**5C),** whereas IMs, monocytes, and PMN showed minimal or no expansion **(Fig.**LJ**S7B).** Changes at 24 h were statistically non-significant **(Fig.**LJ**5C and Fig.**LJ**S7A)**, indicating a delayed cumulative therapeutic effect. LSFM imaging confirmed this robust expansion of the AM pool **(Fig. 5D, E; video S4 vs. S5; Fig. S7C)**, which was driven by local proliferation as evidenced by elevated Ki67 expression specifically in AMs (**Fig. 5F and Fig. S7D**).

We next evaluated fungal uptake and clearance at 6 h, 24 h, and 72 h post infection. AMs engaged more robustly in fungal phagocytosis **(Fig. 5G)** than IMs, PMN, and monocytes **(Fig. S7E).** Mirroring our in vitro data, M-CSF did not alter the AM phagocytosis rate, but significantly enhanced AM-mediated fungal killing at the early 6 h post-infection time point **(Fig. 5H)**, leaving other cell types unaffected **(Fig. S7F).** The absence of enhanced killing at 24 h or 72 h suggests that M-CSF primarily primes the immediate AM response, which is critical for early containment.

### M-CSF prevents fungal-induced pulmonary and systemic Inflammation

To evaluate tissue-level protection, we assessed lung integrity and systemic cytokine profiles. At 72 h post-infection (7 days after allo-HCT), M-CSF–treated mice showed a significantly better preservation of lung architecture than PBS controls (**Fig. 5I**), demonstrating reduced inflammation and collateral tissue damage. This local protection was accompanied by decreased serum levels of pro-inflammatory cytokines (IL-1α, IL-17A, IFN-β) and elevated levels of MCP-1, which may support monocyte recruitment for long-term resolution and tissue repair (**Fig. 5J**).

To uncover the molecular drivers of this uncoupled phenotype, enhanced killing with reduced inflammation, we performed RNA-seq profiling of AMs. Upon *A. fumigatus* exposure, PBS-treated control AMs mounted a classic pro-inflammatory signature consisting of 33 differentially expressed genes (DEGs), including 10 cytokines and chemokines (**Fig. S8A, B, C, Table S1**), the top five of which were validated by qPCR (**Fig. S8D**). Conversely, M-CSF-primed AMs remained transcriptionally quiescent, exhibiting only five DEGs upon fungal encounter (**Fig. S8A, B; Table S2)**. The top two DEGs, validated by qPCR (**Fig. S8E**), were predominantly linked to post-transcriptional modifications and iron homeostasis rather than inflammatory cascades, reinforcing the histological and systemic evidence of blunted inflammation.

### AM-specific reconstitution dictates post-transplant recovery

To investigate the impact of M-CSF on immune cell recovery, we profiled host- and donor-derived pulmonary immune cell compartments at 24 h and 72 h post-infection **(Fig. 6A)**. Neutrophil reconstitution was absent at 24 h post-infection across all conditions, whereas AMs showed a modest, early increase in host-derived populations following M-CSF administration **(Fig. 6B)**.

**Fig. 6:**
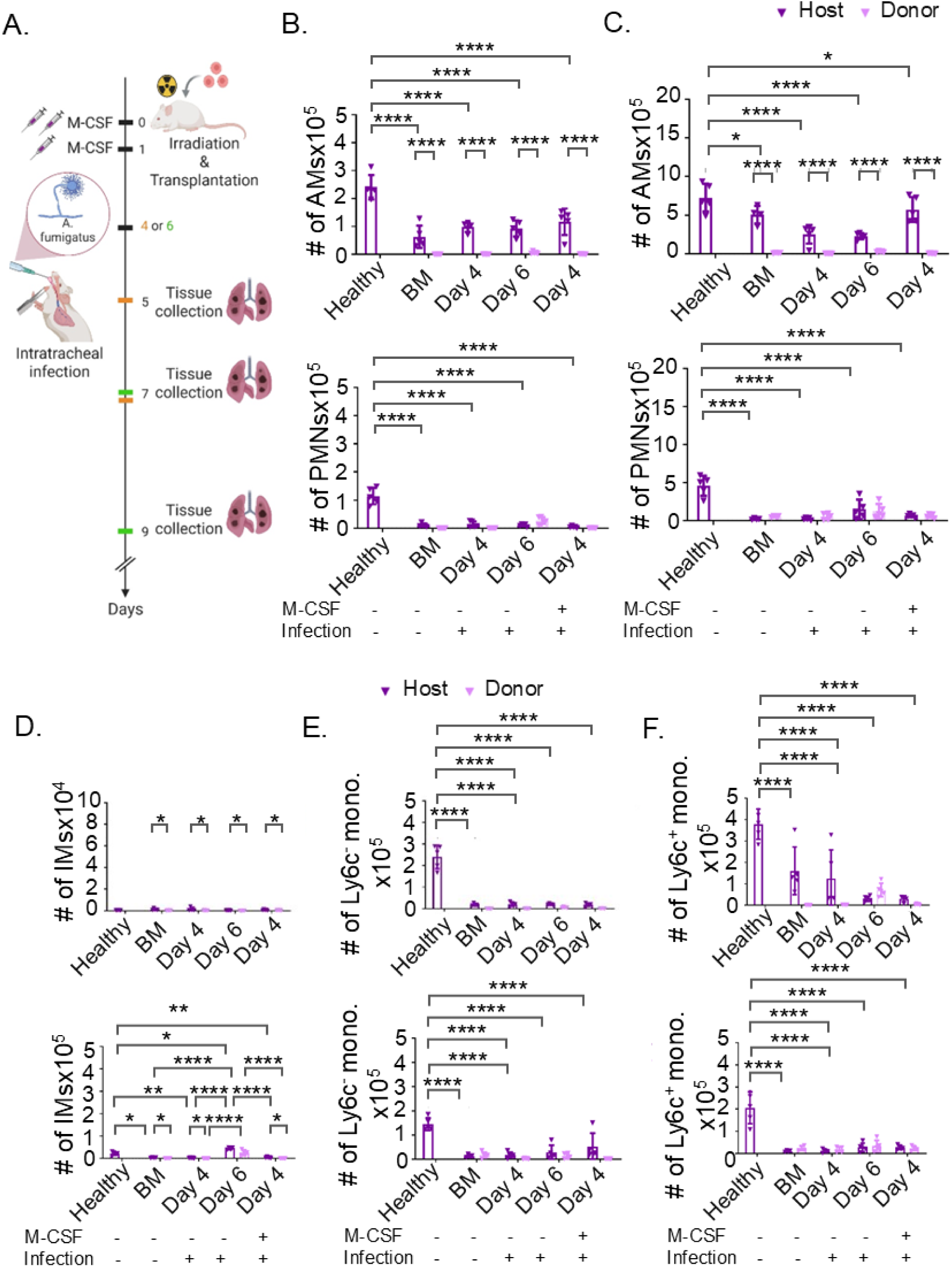
The effect of exogenous M-CSF on pulmonary immune cell recovery. A Experimental scheme for intratracheal *A. fumigatus* infection with a dose of 5×10^4^ ATCC46645 spores per mouse +/- in vivo M-CSF treatment after irradiation with a dose of 8 Gy and allo-HCT of 5×10^6^ BM cells. We injected M-CSF intravenously in 3 doses with a single dose of 4×10^4^ U/120 µl sterile H_2_O at −1, 5, 20 h of allo-HCT. Lungs were collected on day 5 (24 h post infection) and day 7 (72 h post infection) for mice infected at day 4 after allo-HCT, and on day 6 (24 h post infection) and day 9 (72 h post infection) for mice infected at day 6 after allo-HCT. **B, C** Flow cytometry quantification of AMs (B, C upper panel) and neutrophils (B, C lower panel) absolute numbers in the lungs at 24 h (B) and 72 h (C) after *A. fumigatus* infection. n = 5 per group. **D, E, F** Flow cytometry quantification of IMs (D), Ly6c^-^monocytes (E), and Ly6c^+^ monocytes (F) absolute numbers in the lungs at 24 h (upper panel) and 72 h (lower panel) after *A. fumigatus* infection. n = 5 per group. Values were displayed as mean ± SD. Statistical significance was determined by TWO-way ANOVA. (ns, non-significant; *, P ≤ 0.05, **, *p* ≤ 0.01, ***, *p* ≤ 0.001, ****, *p* ≤ 0.0001).

By 72 h post-infection in the day 4 challenge-group, M-CSF treatment significantly expanded AM numbers without altering neutrophil kinetics; however, only a minor recovery was observed in mice challenged at day 6 post-allo-HCT (day 9) **(Fig. 6C)**. In contrast, interstitial macrophages (IMs) and monocytes failed to show significant recovery in either donor- or host-derived compartments at 24 h or 72 h, irrespective of M-CSF treatment or infection timing (**Fig. 6D–F).**

Collectively, these data demonstrate that M-CSF-mediated immune restoration is highly specific to the AM compartment. However, whether this selective expansion and activation of AMs is the sole driver of host survival, or if alternative cellular cross-talk is required, remained to be definitively determined.

### AM depletion abolishes M-CSF-mediated protection against *A. fumigatu in***ifection**

To confirm that M-CSF confers protection against *A. fumigatus* exclusively by enhancing the function of tissue-resident AM function, we locally depleted AMs using a single intratracheal administration of clodronate liposomes 24 h prior to fungal challenge (**Fig. 7A**). Strikingly, AM depletion completely abolished the therapeutic benefit of M-CSF. Regardless of whether they received exogenous M-CSF, AM-depleted mice rapidly succumbed to the day 4 post-allo-HCT infection and required euthanasia within 72 h (**Fig. 7B**). In sharp contrast, non-depleted mice treated with M-CSF consistently maintained a 50-80% survival rate (**Fig. 7B**), recapitulating our earlier findings (**Fig. 5B**). Flow cytometry of whole-lung digests at 72 h post-infection confirmed that the intratracheal clodronate liposomes selectively eliminated the AM pool, whereas pulmonary monocyte and PMN counts remained entirely unaffected (**Fig. 7C, D**). These loss-of-function experiments definitively demonstrate that the protective efficacy of M-CSF against early post-transplant *A. fumigatus* infection operates through an AM-dependent mechanism, underscoring the pivotal role of these tissue-resident cells as the primary gatekeepers of the immunocompromised lung.

**Figure 7:**
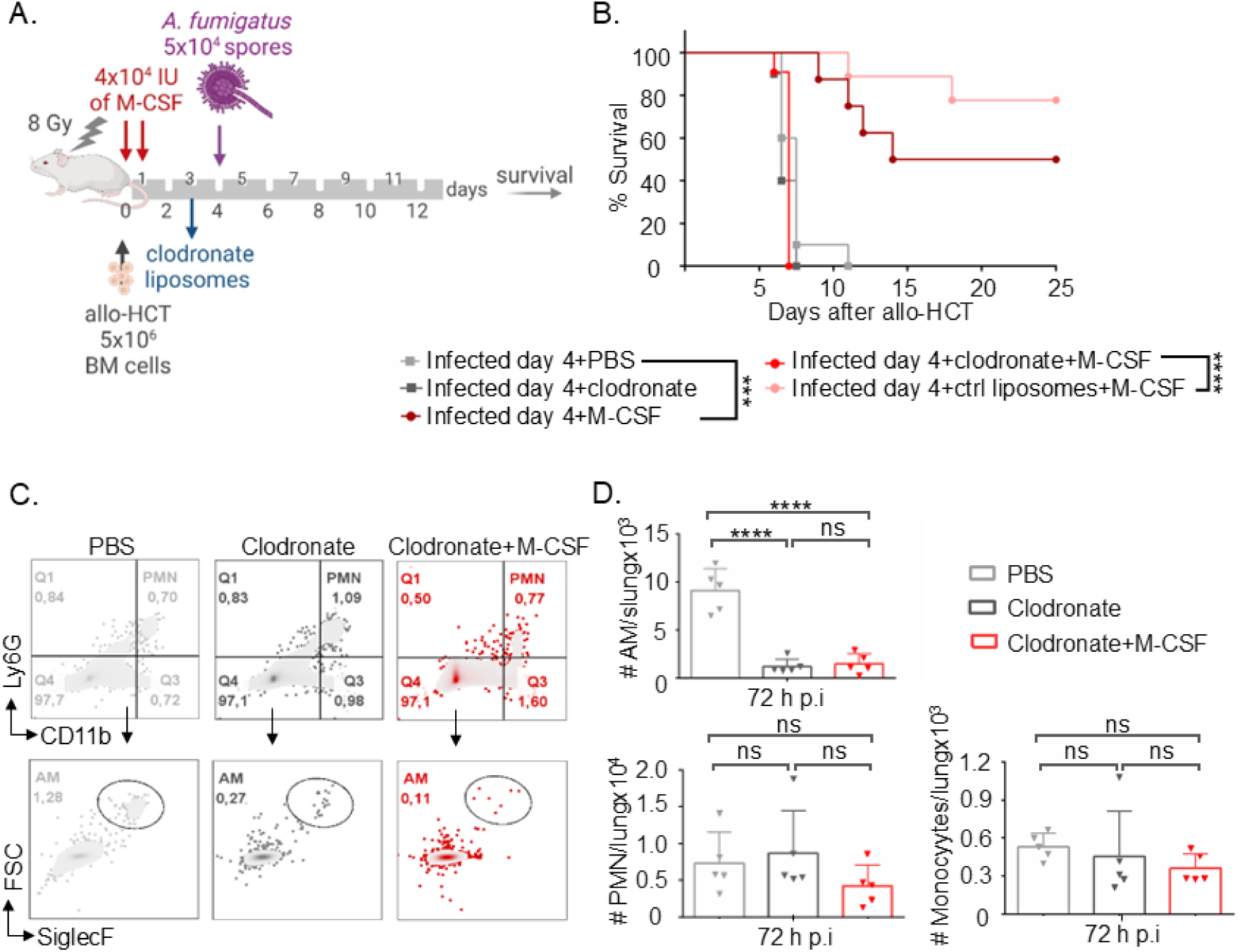
In vivo M-CSF treatment of *A. fumigatus*-infected mice with depleted AMs. (**A**) Experimental scheme for AM depletion and in vivo M-CSF treatment of mice prior to intratracheal *A. fumigatus* infection. We infected the mice with a dose of 5×10^4^ ATCC46645 spores per mouse at day 4 after irradiation and allo-HCT. AMs were depleted by intratracheal administration of 50 μl of clodronate liposomes 24 h before infection. (**B**) Survival of in vivo M-CSF-treated, AM-depleted and infected mice compared to M-CSF-treated controls, AMs-depleted controls and PBS-treated controls. Statistical significance was determined by Mantel-Cox test. *n* = 8-11 per group. (**C, D**) Representative flow cytometry plots for AM depletion (C) Flow cytometry quantification of AMs, PMN, and monocytes numbers (D) at 72 h after *A. fumigatus* infection +/- M-CSF treatment and +/- AM depletion. *n* = 5 per group. Statistical significance was determined by ordinary one-way ANOVA with Holm-Sidak’s multiple comparisons test. Values were displayed as mean ± SD. (ns, not significant, ****, *p* ≤ 0.0001).

## Discussion

Host susceptibility to pulmonary *A. fumigatus* infections relies heavly on innate immune coordination^9^. While immunocompetent hosts rapidly clear inhaled conidia, immunocompromised patients, particular those undergoing allo-HCT, remain exceptionally vulnerable to IPA^7,57,58^. This vulnerability has traditionally been attributed to delayed systemic immune reconstitution, pattern recognition receptor (PRRs) polymorphisms (e TLR-4^59–61^ and Dectin-1^62^) or defects in antigen presentation, such as CCR7 deficiency on DCs^63^. This view, however, overlooks the contribution of tissue-resident immune compartments. We demonstrate that the functional state of lung-resident AMs, rather than systemic immune numbers, is a key determinant of antifungal protection post-allo-HCT, shifting the focus toward a previously underappreciated cellular niche and opening new avenues for targeted immunotherapeutic interventions.

Traditional infection models often employ supra-physiological doses of conidia or hyper-virulent fungal strains that provoke intense but transient lung inflammation ^24,27^ and trigger neutrophil-driven clearance mechanisms^25^. While informative, these models fail to recapitulate the unique immune vulnerabilities of allo-HCT recipients and often rely on depletion of specific immune subsets within an immunocompetent immune system to infer their roles in host defense^24^. To overcome these limitations, we established a clinically relevant murine model incorporating host conditioning through myeloablative total body irradiation, allo-HCT, and inoculation with a physiologically relevant dose of *A. fumigatus* (∼5L×L10L conidia/mouse) instead of megadose infection systems (10L–10L conidia/mouse). In this setting, tissue-resident AMs emerged as the predominant and functionally essential immune population. In contrast to neutrophils and peripheral monocytes, AMs retained potent phagocytic and fungicidal capacity. Their depletion resulted in near-universal mortality, pinpointing AMs as indispensable early sentinels. This aligns with a previous study reporting persistent fungal burden in AM-deficient lungs despite substantial neutrophil infiltration, even in immunocompetent models using high-dose infections^64^. Our work extends these findings by demonstrating the non-redundant role of AMs in antifungal defense during the critical early window following allo-HCT.

Our findings challenge the neutrophil-centric pardigm of antifungal defense by revealing a critical role for AMs in early *A. fumigatus* control, particularly in the context of allo-HCT^65,66^. Our infection model contradicts previous reports where neutrophils compensated for AM loss^24,67^. We demonstrate, however, that under immune-suppressed conditions and with physiological fungal burdens, there is no compensation from neutrophils. We show that the timing of infection relative to immune reconstitution is a key determinant of outcome: survival was near-complete when infection occurred on day 6 after allo-HCT but plummeted when mice were infected two days earlier. This underscores the fragile, dynamic nature of early alveolar immunity, where AMs serve as essential sentinels. Our profiling of pulmonary immune cell composition demonstrates the predominance of AMs during this period, highlighting their central role while other effector immune populations recover only gradually over time.

Importantly, we identify M-CSF as a potent enhancer of AM antifungal activity. We find that M-CSF selectively induces AM proliferation, cytoskeletal remodeling, and a transition from sessile to motile behavior. This increase in migratory behavior enables more efficient alveolar surveillance, as supported by previous studies^27,28^. Unlike sessile AMs, which rely on connexin-43-(Cx43)-mediated interactions with AEC^28^ or passive AEC-driven conidial displacement, motile AMs actively patrol the alveolar niche via LFA-1-mediated adhesion^27^. Notably, this phenotype was tissue-specific and not observed in peritoneal macrophages.

In our hands, M-CSF-treatment of AMs decoupled phagocytosis from killing efficacy. We observed that while fungal uptake rates remained unchanged, phagolysosomal acidification and cathepsin B activity were significantly increased. This contrasts with an earlier report using rabbit AMs, which observed that M-CSF enhanced phagocytosis^68^. However, that study lacked AECs in the co-culture assays. Importantly, using an AEC–AM–

*A. fumigatus* co-culture system we observed that AEC internalize spores, delay germination and promote AM motility, potentially buffering early fungal growth while facilitating AM activation and recruitment. Notably, the full antifungal potential of M-CSF–stimulated AMs only became apparent in the presence of epithelial cells, underscoring the importance of tissue context and intercellular communication.

M-CSF has traditionally been studied in the context of monocyte-derived macrophages, where it was shown to enhance antifungal activity in monocytes and peripheral macrophages by increasing phagocytosis and triggering hyphal damage^68,69,70^. Moreover, M-CSF was associated with protection against *A. fumigatus* infection post-allo-HCT^71^, but it was not clear which specific cell types are mediating this effect. Here, we demonstrate that M-CSF directly affects lung-resident AMs and promotes their proliferation and motility. These effects are not replicated by IL-34, although both cytokines engage the same receptor^72,73^. These distinct outcomes might be due to differences in receptor binding sites, signaling intensity and activation kinetics. IL-34 is known to drive pro-inflammatory programs^74^ and our morphological and metabolic profiling also revealed that IL-34 preferentially promotes a rounded, inflammatory phenotype and lipid mediator release, especially during fungal co-culture. In contrast, M-CSF supports an elongated morphology and restrained inflammatory response – traits that are particularly advantageous in allo-HCT, where rapid fungal clearance and minimizing excessive inflammation and tissue damage are key to avoiding immunopathology. Our molecular and lipidomic data suggest that the subdued inflammatory milieu seen with M-CSF is reflective of a cytokine program that promotes pathogen clearance without collateral tissue damage. This anti-inflammatory, pro-resolving profile may also explain the improved survival and preserved alveolar architecture observed in vivo following M-CSF treatment. Critically, selective AM depletion abolished these benefits, reinforcing the conclusion that M-CSF acts via AM stimulation to confer both immune protection and tissue preservation.

While our model recapitulates key features of immune suppression post-allo-HCT, it does not reflect the full clinical complexity of transplant recipients, including microbial dysbiosis, viral reactivations, or alloimmune phenomena. Moreover, although our co-culture system incorporates key epithelial–AM interactions, it lacks the full 3D architecture and mechanical dynamics of the alveolar space. Future work using single-cell transcriptomics, spatial mapping, or intravital imaging will be essential to define how M-CSF shapes AM identity and function within the native lung environment.

Nevertheless, our findings carry immediate translational relevance. AM function is currently not assessed in clinical allo-HCT protocols, yet our study identifies AMs as central determinants of survival during early infection. The identical response of human AMs to M-CSF highlights its potential as a promising immunotherapeutic candidate. Although M-CSF is not FDA-approved yet, its early clinical signals, such as improved outcomes in *Candida*-infected transplant patients in a Phase I/II study^75^, and enhanced survival in transplant recipients with invasive fungal diseases when used alongside conventional antifungal therapy^76^, highlight its promise. Successful clinical translation will require rigorous pharmacokinetic and safety profiling of M-CSF in immunocompromised patients. Targeting tissue-resident immunity via M-CSF may offer a novel strategy to mitigate IPA risk in allo-HCT recipients, particularly during the early reconstitution window when conventional blood-based markers fail to capture critical deficits in lung immunity.

## Materials and methods

### Mice

All experiments were performed using female 10- to 20-week-old mice. BALB/cAnNRj mice were either obtained from Janvier Labs (Paris, France) or bred in-house at IMIB (Institute for Molecular Infection Biology, University of Würzburg, Germany). C57BL/6 mice were bred in-house at ZEMM (Center for Experimental Molecular Medicine, Würzburg University Hospital, Germany). Mice were randomly assigned to experimental groups and housed no more than five per cage with Aspen chip 2 bedding. Mice were maintained under a 12-hour light and 12-hour dark cycle at 20–22°C. Water and food were provided freely, and mice were fed high-calorie diet when needed. All experimental procedures complied with German regulations for animal experimentation and received approval from the authorities of lower Franconia under permit numbers (55.2-2531.01-86-13, 55.2-2532-2-403, and 55.2.2-2532-2-1627).

### Human lung specimens

Fresh human lung biopsies with a size range of 2–4 cm^3^ were provided by Würzburg University Hospital and were processed immediately after lung segmentectomy or lobectomy. The lung biopsies used in this study were from male and female patients with an age range of 52–82 years. The patients suffered from underlying malignancies with either a smoking or non-smoking history. The lung biopsies of heavy smokers were excluded from the study. Ethical approval to use lung biopsies was granted by the authorities of lower Franconia under permit number 34/22 and informed consent was obtained from all participants after the nature and possible consequences of the studies were explained.

### Pathogens

For phagocytosis assays, *Aspergillus fumigatus* FLARE conidia were used (generously provided by Prof. Tobias Hohl^44^). Other experiments utilized the ATCC46645 strain. FLARE conidia were surface labeled with Alexa Fluor 633 (AF633) and used for tracking fungal fate. Fungal viability was assessed based on the dsRed signal of FLARE conidia, which diminishes upon death. *A. fumigatus* was grown on *Aspergillus* minimal medium agar plates^77^ at 37°C and 5% CO_2_ and freshly harvested before the experiments.

The *Staphylococcus aureus* USA300_JE2 RFP-reporter strain was constructed using an RFP-expressing plasmid, kindly provided by Martin Fraunholz. An infection stock from *S. aureus* USA300_JE2 expressing RFP was grown overnight at 37 °C in BHI medium supplemented with chloramphenicol at 10 µg/ml final concentration and stored as infection stocks at −80 °C. Bacterial cells from infection stocks were washed twice using sterile saline and resuspended to the desired CFU/ml (based on a pre-constructed standard calibration curve). Bacterial phagocytosis relied on the RFP signal, while bacterial killing was determined after performing extracellular bacterial killing for 1 h using gentamicin. Lysed AMs were plated on LB agar plates and engulfed bacterial CFUs growing overnight at 37°C and 5% CO_2_ were counted. Specific information on the pathogens used in the study can be found in Supplementary Table 1.

### Bronchoalveolar lavage (BAL)

Mice were euthanized by CO_2_ overdose, fixed on the dorsal side and sprayed with 70% ethanol. The trachea was located by cutting through the skin covering the neck and dissecting away the surrounding tissue (salivary glands, muscles and larynx, etc.). Through a small incision in the trachea, a catheter (Vasofix® Safety 1,30 x 45 mm G 18, B.Braun) connected to 1 ml syringe was inserted and the lungs were flushed with 1ml 0.6 mM EDTA phosphate-buffered saline (PBS) for 8-10 times with flushing 2 to 3 times in each step. Recovered cells were suspended in RPMI 1640 supplemented with 10% fetal calf serum (FCS) and 1% penicillin G/streptomycin (pen/strep), yielding approximately 4–5 x 10^5^ AMs per mouse with ∼85% purity.

### Isolation of human AMs (huAMs)

Fresh human lung biopsies from Würzburg University Hospital were processed immediately as previously described^78,79^. Briefy, lung tissue was dissected and blood cells lysed with FACS lysis buffer (9 mM NH_4_Cl, 1 mM KHCO_3,_ and 0.037 mM EDTA in ddH_2_O) for 10 min. After washing, cell pellets were resuspended in RPMI 10% fetal bovine serum (FBS), 1% pen/strep and seeded overnight in 75T flask at 37 °C and 5% CO_2_. After 24 h, cells were washed twice with warm PBS to remove non-adherent cells. Adherent huAMs were then harvested using accutase for 15–20 min at 37 °C and 5% CO_2_, washed with warm PBS and resuspended in RPMI 10% FBS, 1% pen/strep for cytokine stimulation.

### Culture of alveolar epithelial cells (AECs)

AEC were cultured in T25 or T75 cell culture flasks pre-coated with gelatin solution. Murine and human AEC were maintained by culture in complete epithelial cell medium with kit and human complete epithelial cell medium with kit, respectively and incubation at 37°C with 5% CO_2_ where the medium was changed daily. For AECs recovery, cells were incubated with 3-5 ml pre-warmed 0.05% trypsin/EDTA for 5 min at 37°C with 5% CO_2_. The reaction with trypsin was stopped by adding 8–10 ml of fresh AEC medium. Afterwards, cells were collected and centrifuged at 120 g for 5 min, then the pellet was resuspended in the respective AEC medium.

### In vitro cytokine stimulation of murine and human AMs

AMs were cultured in RPMI medium with 10% FCS and 1% pen/strep in 24-well plates with a density of 3-8×10^5^ cells per ml. Human and murine AMs were stimulated with M-CSF, IL-34, GM-CSF or G-CSF for 24 h and 48 h, with concentrations of 300 IU/ml, 6000 IU/ml, 6000 IU/ml, and 6000 IU/ml, respectively. Specific information on the cytokines used in the study can be found in Supplementary Table 1,

### Time lapse imaging of fungal phagocytosis by AMs

Cytokine-pretreated murine AMs or human AMs were stained with 1:400 anti-mouse SiglecF or 1:20 anti-human CD206 and 1:20 anti-human CD169, respectively, for 30 min at 4°C. 1×10^5^ AMs were cultured with 1:3 tdTomato or ATCC46645 spores in 30 µl of RPMI 1640 with 10%FCS and 1% pen/strep and seeded in VI 0.4 µ-Slide (ibidi), which was pre-seeded with 5 ×10^4^ alveolar epithelial cells (AECs) in complete AEC medium for 6 h at 37°C and 5% CO_2._ Then, 60 µl of fresh culture medium was added on both sides of each chamber in the slide before imaging. Fungal phagocytosis was followed for up to 15 h with an increment of 2 min (AMs) and 3 min (huAMs) between each acquired image at 37°C with confocal microscopy (LSM 780, ZEISS, Germany). AM and huAM time lapse movie have 300 and 450 frames, respectively, with a pixel width and height of 0.415 µm. AM migration was quantitatively analyzed by tracking each AMs manually with ImageJ. AM speed, migration behavior, and directionality were quantified with MATLAB. To measure cell migration speed and the distance travelled, mean squared displacement (MSD) was assessed as described previously^80^. Diffusion co-efficient (D) was calculated from the slope of linear fit to MSD curve. Both MSD and turning angle distribution were combined to determine the type of AM locomotion.

### In vitro phagocytosis assay

Cytokine-stimulated AMs were seeded on pre-incubated AEC (2.5 ×10_4_ cells/well in 96-well plate for 2 h at 37°C and 5% CO_2_) with a density of 1 ×10^4^-1×10^5^ cells/100 µl. FLARE conidia, *Staphylococcus aureus* were used in MOI (multiplicity of infection) of 1:1 or 1:5 (AMs: pathogen) and incubated with AMs for 1 h or 6 h at 37°C and 5% CO_2_ in 200 µl RPMI 1640 with 10% FCS. After co-culture, plates were centrifuged, washed with PBS. For murine AMs, cells were blocked for 10 min with 5% NRS at 4°C. AMs were stained with 1:100 Zombie Aqua^TM^ Fixable Viability and identified using 1:400 anti-mouse SiglecF for 30 min at 4°C. For huAMs, cells were stained in human cell wash buffer and identified using 1:20 anti-human CD206, and 1:20 anti-human CD169 for 30 min at 4°C. Before measuring plates were fixed with 4% paraformaldehyde (PFA) for 30 min at RT.

### In vitro reactive oxygen species (ROS) production

Cytokine-stimulated AMs were seeded on pre-incubated AEC (2.5 ×10^4^ cells/well in 96-well plate for 2 h at 37°C and 5% CO_2_) with a density of 1-1.5×10^5^ cells/100 µl. ATCC46645 conidia, were used in MOI of 1:5 (AMs: *A. fumigatus*) and incubated with AMs for 6 h at 37°C and 5% CO_2_ in 200 µl RPMI 1640 with 10% FCS and 1% pen/strep. After co-culture, plates were centrifuged, washed with PBS and blocked for 10 min with 5% NRS at 4°C. AMs were stained with 1:100 Zombie Aqua^TM^ Fixable Viability and identified using 1:400 anti-mouse SiglecF for 30 min at 4°C. To stain for ROS, cells were incubated in 100 µl of 20 µM of DCFDA ROS reagent (2′,7′-Dichlorodihydrofluorescein diacetate) for 30 min at 4°C. Before measuring plates were fixed with 4% paraformaldehyde (PFA) for 30 min at RT.

### In vitro analysis of lipid mediators

Cytokine-stimulated AMs were seeded on pre-incubated AEC (2.5 ×10^4^ cells/well in 96-well plate for 2 h at 37°C and 5% CO_2_) with a density of 1-1.5×10^5^ cells/100 µl. ATCC46645 conidia, were used in MOI of 1:5 (AMs: *A. fumigatus*) and incubated with AMs for 6 h at 37°C and 5% CO_2_ in 200 µl RPMI 1640 with 10% FCS and 1% pen/strep. After co-culture, plates were centrifuged, and 150 µl of the supernatant were collected for the analysis of lipid mediator production. The supernatant was snap frozen in liquid nitrogen before transfer to – 80°C for long-term storage. Sample preparation was conducted by adapting published criteria^81^. In brief, samples were kept at −20 °C for 60 min to allow protein precipitation. After centrifugation (1200 g, 4 °C, 10 min) 9 mL acidified H_2_O was added (final pH = 3.5) and samples were subjected to solid phase extraction. Solid phase cartridges (Sep-Pak^®^ Vac 6cc 500 mg/ 6 mL C18; Waters, Milford, MA) were equilibrated with 6 mL methanol and 2 mL H_2_O before samples were loaded onto columns. After washing with 6 mL H_2_O and additional

6 mL *n*-hexane, LM were eluted with 6 mL methyl formate. Finally, the samples were brought to dryness using an evaporation system (TurboVap LV, Biotage, Uppsala, Sweden) and resuspended in 150 µL methanol-water (50/50, v/v) for UPLC-MS/MS automated injections. LM profiling was analyzed with an Acquity™ UPLC system (Waters, Milford, MA, USA) and a QTRAP 5500 Mass Spectrometer (ABSciex, Darmstadt, Germany) equipped with a Turbo V™ Source and electrospray ionization. LM were eluted using an ACQUITY UPLC^®^ BEH C18 column (1.7 µm, 2.1 × 100 mm; Waters, Eschborn, Germany) at 50 °C with a flow rate of 0.3Lml/min and a mobile phase consisting of methanol-water-acetic acid of 42:58:0.01 (v/v/v) that was ramped to 86:14:0.01 (v/v/v) over 12.5 min and then to 98:2:0.01 (v/v/v) for 3 min^82^. The QTrap 5500 was operated in negative ionization mode using scheduled multiple reaction monitoring (MRM) coupled with information-dependent acquisition. The scheduled MRM window was 60 sec, optimized LM parameters were adopted^83^, and the curtain gas pressure was set to 35Lpsi. The retention time and at least six diagnostic ions for each LM were confirmed by means of an external standard (Cayman Chemical/Biomol GmbH, Hamburg, Germany). Quantification was achieved by calibration curves for each LM. Linear calibration curves were obtained for each LM and gave r2 values of 0.998 or higher (for fatty acids 0.95 or higher). Additionally, the limit of detection for each targeted LM was determined.

### Fungal burden

Mice were euthanized by CO_₂_ inhalation, and one-half of each lung was aseptically collected, weighed, and placed in ice-cold PBS. Lung tissue was homogenized by passing it through a sterile 70-µm cell strainer using the plunger of a 5-mL syringe. The strainer was pre-wetted with 1 mL PBS and rinsed with an additional 4 mL PBS to recover the homogenate. The undiluted homogenate was used for fungal burden determination. A 100-µL aliquot of the homogenate was spread onto Aspergillus minimal medium (AMM) agar plates in triplicate and incubated at 37 °C for 48 h. Colonies were counted, and fungal burden was expressed as colony-forming units (CFU) per gram of lung tissue.

### RNA sequencing

For in vitro infection assays, AMs (1×10^4^ per condition) were stimulated with either M-CSF or PBS for 48 h in cell culture medium at 37°C and 5% CO_2_. AMs were arranged in such a way that each condition had biological triplicates. After cytokine treatment, cells were washed with PBS, then *A. fumigatus* ATCC46645 spores were added to the corresponding wells in MOI of 1:5 (AMs: *A. fumigatus*) and incubated with AMs in RPMI medium with 10% FCS and 1% pen/strep for 6 h at 37°C and 5% CO_2_. In addition, PBS-treated AMs without conidia were used as baseline controls. To isolate RNA, AMs were washed two times with ice-cold PBS and re-suspended in 50 µl lysis buffer (95 µl of 10X lysis buffer + 5 µl RNase inhibitor + 900 µl H_2_O) from the SMART-Seq® v4 Ultra® Low Input RNA Kit. AM lysates were transferred to RNase-free PCR tubes and stored at −80°C until further analysis. Library preparation was performed using the SMART-Seq® v4 Ultra® Low Input RNA Kit according to manufacturer’s instructions. The PCR amplification was performed using 12 PCR cycles. Libraries were quantified by Qubit^TM^ dsDNA HS Assay Kit and quality was checked using 2100 Bioanalyzer with High Sensitivity DNA kit (Agilent Technologies). 0.5 ng of each library was subjected to a tagmentation-based protocol (Nextera XT, Illumina) using a quarter of the recommended reagent volumes. Libraries were quantified again by Qubit^TM^ dsDNA HS Assay Kit and quality was checked using 2100 Bioanalyzer with High Sensitivity DNA kit before pooling. Sequencing of pooled libraries, spiked with 1% PhiX control library, was performed in single-end mode with 75 nt read length on the NextSeq 500 platform (Illumina) with a High Output Kit v2.5. Demultiplexed FASTQ files were generated with bcl2fastq2 v2.20.0.422 (Illumina). Following sequencing and genome mapping using HISAT2, a list of differentially expressed genes (DEGs) was detected according to the EdgeR algorithm and based on Median Ratio Normalization (MRN) with a log2 fold change cut off value of 1 for upregulated genes and −1 for downregulated genes. The data discussed in this publication have been deposited in NCBI’s Gene Expression Omnibus (Edgar *et al*., 2002) and are accessible through GEO Series accession number GSE291042 (https://www.ncbi.nlm.nih.gov/geo/query/acc.cgi?acc=GSE291042).

qRT-PCR was performed for representative DEGs to confirm the results of dual RNA-seq. Briefly, using 24 well plate, 0.8 - 3 ×10^6^ AMs were stimulated with either M-CSF, Il-34 or PBS 48 h in RPMI medium with 10% FCS and 1% pen/strep at 37°C and 5% CO_2_. AMs were arranged in such a way that each condition has three biological triplicates. After cytokine stimulation, cells were washed with PBS then ATCC46645 conidia were added to the corresponding wells with MOI of 1:5 and incubated with AMs in RPMI medium with 10% FCS and 1% pen/strep for 6 h at 37°C and 5% CO_2_. After 6 h, the plate was centrifuged, and RNA was isolated from AMs and *A. fumigatus* conidia using commercially available RiboPure^TM^-yeast RNA purification kit (Thermo Fisher). RNA concentration was determined using Nanodrop 2000 spectrophotometer (Thermo Fisher Scientific). Subsequently, RNA was reverse–transcribed reverse-transcribed to cDNA using the iScript^TM^ reverse transcription supermix (Bio-Rad) according to manufacturer’s instructions.

qRT-PCR was performed on a CFX Connect Real-Time PCR System (Bio-Rad). 10 ng cDNA was used per PCR reaction. To prepare a qPCR reaction (in total 10 µl), 1 µl of cDNA was mixed with 5 µl SsoAdvanced universal SYBR Green Supermix (Bio-Rad), 0.5 µl of forward and reverse primers, respectively (final concentration: 0.5 mM), and 3 µl nuclease-free water (Thermo Fisher). Primer pairs (Sigma Aldrich) used for qRT-PCR are listed in Table 1. PCR reactions were done in technical triplicates. Additionally, for each DEG verified by qRT-PCR, a non-reverse transcribed control (reaction mix with 10 ng RNA) per biological replicate and a water control were included. PCR cycling routine used for all reactions was set up as follows: 95°C, 30 sec (polymerase activation); 40 cycles of 5 sec at 95°C (denaturation), 20 sec at 60°C (annealing/extension and plate read). Melt curve analysis was done under the following condition: 65 to 95°C, 0.5°C increments at 5 sec/step. In total 17 genes were checked with RT-qPCR for the comparison between different treatment conditions. RPL13A^84^ was used as reference housekeeping genes of AMs, for quantitative analyses using the comparative threshold cycle method (2^-ΔΔCT^ method)^85^.

For CSFR1 expression analysis, lungs were harvested and one lung from each mouse was processed for isolation of immune or non-immune cells, respectively. To isolate immune cells, lungs were digested in Liberase (50 µg/ml; Roche) and DNase I (25 µg/ml; Sigma Aldrich) in Iscove’s Modified Dulbecco’s medium (IMDM) medium (Gibco) for 30 min at 37 °C and mechanically dissociated using the gentleMACS Octo Dissociator with Heaters (Miltenyi Biotec). To isolate non-immune cells, lungs were digested in RPMI 1640 medium (Gibco) containing DNase I (0.1 mg/ml), dispase II (0.8 mg/ml; Sigma-Aldrich), collagenase P (0.2 mg/ml; Roche) for 45 min at 37 °C under constant rotation. The reaction was stopped at 45 min with 3 ml heat-inactivated FBS (Gibco). Cells were treated with red cell lysis buffer (0.15LmM NH_4_Cl, 10LmM NACO_3_, 1 mM EDTA), washed with medium and filtered through a 70 µm cell strainer. Cell numbers and viability were quantified on the LUNA-FL™ Dual Fluorescence Cell Counter (Logos Biosystems) using Acridine Orange/Propidium Iodide Stain (Logos Biosystems). For the non-immune cell population, CD45 positive cells were removed using the EasySep Mouse CD45 positive selection kit (Stem Cell technologies). Equal numbers of immune and non-immune cells (5×10^5^ cells per population) from the lung of each mouse (n = 3 per treatment group) were pooled and fixed with the Evercode Cell Fixation kit v3 (Parse Biosciences). Single-cell RNA sequencing libraries were prepared using the Evercode WT Kit v3 (Parse Biosciences) according to the manufacturer’s instructions. Libraries were sequenced on an Illumina NovaSeq X Plus platform using a 10B Flow cell. Data was analysed using the Parse Biosciences Trailmaker platform. Sequencing data will be deposited in the Gene Expression Omnibus (GEO) database upon publication.

### Structured illumination microscopy (SIM)

AMs (seeded on PDL-coated glass cover slips) were permeabilized for 1-2 min in 0.3% glutaraldehyde and 0.25% Triton-X-100 in cytoskeleton buffer (CB) (10 mM MES pH 6.1, 150 mM NaCl, 5 mM EGTA, 5 mM Glucose, 5 mM MgCl2) then fixed for 10 min in 2% glutaraldehyde in CB at 4°C. After washing with PBS and blocking for 30 min in 5% bovine serum albumin (BSA)/PBS, cell cytoskeleton was stained with 1:100 anti ß-tubulin primary antibody in 5% BSA/PBS for 60 min. After washing twice with 0.1% Tween20/PBS for 5 min, 1:200 of secondary antibody, goat anti-mouse F(ab)2-fragments were used for 45 min. After washing, cells were fixed with 4% formaldehyde/PBS for 5 min, washed with PBS then stained with 1:100 anti-phalloidin in PBS overnight and washed with PBS immediately before imaging. Imaging was performed using SIM Zeiss Elyra S.1.

### Scanning electron microscopy (SEM)

BAL-isolated AMs, which were not co-cultured with *A. fumigatus* ATCC44546, were directly added on top of 12 mm-coverslips in a 24-well plate and stimulated with M-CSF or IL-34 as previously described. AMs designated for co-culture with *A. fumigatus* ATCC46645 were stimulated in a 96-well round-bottom plate for 48 h. Prior to co-culture, 2 x 10^5^ AEC were seeded on 12 mm-coverslips for 4 h for them to adhere. Subsequently, 1 x 10^5^ pre-stimulated AMs were added together with *A. fumigatus* spores in a MOI 1:5 and incubated at 37 °C with 5 % CO_2_ for 1 h, after which the plate was centrifuged for 5 min at 1200 rpm. The supernatant was carefully removed, and cells were fixed with 6.25% glutaraldehyde in phosphate buffer overnight. Samples were further processes for SEM imaging as described in Grob et al., 2020^86^. Imaging was performed using SEM JEOL JSM-7500F.

### Detection of lysosome and lysosomal cathepsin B

Human and murine AM were seeded on top of 10 mm-coverslips in a 48-well plate and stimulated with M-CSF or IL-34 as previously described. After stimulation, cells were washed with PBS and *A. fumigatus* ATCC46645 spores were added in MOI 1:5 and incubated for 1 h at 37 °C and 5% CO_2_. After fungal co-culture, cells were washed with PBS and incubated with Acridine Orange (1:1000 dilution of stock solution in PBS) for 30 min at 37 °C and 5%

CO_2_ or with Magic Red solution (1× final concentration) for 50 min at 37 °C and 5% CO_2_. Nuclei were stained using Hoechst 33342 solution (1:200 dilution) for 10 min at 37 °C. After staining, cover slips were mounted on slides and immediately imaged using confocal microscopy (LSM 780, ZEISS) and analyzed using FIJI image analysis software. Corrected total cell fluorescence (CTCF) of Magic Red (MR) and Acridine Orange (AO) was quantified using the following formula: CTCF = Integrated Density – (Area of selected cell X Mean luorescence of background readings).

### Allo-HCT and *Aspergillus fumigatus* infection

BALB/c recipient mice were myeloablatively irradiated (8 Gy TBI, Mevatron Primus, Siemens, Germany). Within 3 h after irradiation, the mice were transplanted (i.v) with 5×10^6^ bone marrow (BM) cells obtained from untreated C57BL/6 donor mice. For *A. fumigatus* infection, after mice were anesthetized, the tracheal opening is located and a 22G catheter (B.Braun) is inserted into the trachea. Each mouse was intratracheally infected with a dose of 5×10^4^ ATCC 46645 f spores suspended in 50 µl of 0.9% NaCl + 0.005% Tween-20 either at day 4 or day 6 after HCT. The catheter was immediately connected to a ventilator (stroke volume 250 µl, 225 stroke/ min) for 2 min to maintain an even distribution of spores in the mouse lungs.

### In vivo M-CSF treatment

BALB/c recipient mice received M-CSF treatment as three intravenous injections at −1 h, 5 h, 20 h after HCT. M-CSF lyophilized powder was freshly reconstituted with sterile distilled water and injected into the mice in a concentration of 40 µg (120 µl) per each injection.

### Cytometric bead array

To analyze serum cytokine and chemokine levels, 50 µl of blood were collected from tail vein and incubated 30 min at RT to allow serum separation. Serum was analysed using LEGENDplex^TM^ mutianalyte flow assay kit and according to manufacturer’s protocol.

### Lung histology

Mice were sacrificed and lungs were extracted 72 h after *A. fumigatus* infection. Lungs were fixed for 24 h in 4% PFA then each lobe was cut into smaller parts and stained with haematoxylin and eosin (H&E).

### Flow cytometry and lung preparation

To prepare lung single-cell suspension, the lungs were digested using gentleMACS dissociator (Miltenyi Biotec) and lung dissociation kit according to manufacturer’s protocol. To stain for flow cytometry, dead cells were excluded by staining with Zombie Aqua^TM^ Fixable Viability kit and cells were characterized by antibody staining including anti-CD45.1, anti-F4/80 anti-CD11c, anti-SiglecF, anti-CD11b, anti-Ly6G, anti-Ly6C, MHC II, and CD64. Specific information on the antibodies used in the study can be found in Supplementary Table 1.

For measuring cell proliferation, anti-Ki67^+^ intracellular staining was performed. Flow cytometry was performed with Attune NxT Flow Cytometer (Thermofisher Scientific) with the use of anti-mouse CompBeads (BD) for fluorescence compensation. To analyse the data FlowJo Software version 10 (Tree Star, Ashland, OR) was used.

### LSFM and lung preparation

Before lung extraction, mice were perfused using an ISMATEC Reglo Analog pump (IDEX Health & Science LLC, Oak Harbor, WA, USA). Shortly, the mice were anesthetized by intraperitoneal injection of ketamine (100 mg/kg) and xylazine (10 mg/kg). Surgical tolerance was established by checking the plantar reflex, then the lungs of the mice were exposed and a needle for infusion was inserted into the left ventricle. The right atrium was incised to allow the perfusion solutions to escape while a peristaltic pump is used to pump ∼100 ml of PBS for 2 min and then ∼300 ml of 4% PFA for 8 min through the circulatory system.

To prepare the samples for imaging, lungs were further fixed in 4% PFA for 2 h at RT, then washed with PBS three times for 30 min each. The lungs were incubated in a blocking solution (2% fetal calf serum (FCS), 1% normal rat serum (NRS), 1% bovine serum albumin

(BSA) and 0.3% Triton X-100 in PBS) for 24 h at 4°C. To stain for immune cells, the following antibodies were used; anti-F4/80 (A488), anti-CD11c (A532), anti-SiglecF (Dy755), anti-CD11b (A488 or A647), anti-Ly6G (A750), and anti-Ly6C (A488). For *A. fumigatus* visualization, the monoclonal antibody JF5 (DyLight 655) was used^43^. All antibodies were prepared in a titer of 1:100 in blocking solution and added to the lungs for 48 h at 4°C. Specific information on the antibodies used in the study can be found in Supplementary Table 1. After staining, lungs were washed with PBS three times for 30 min each and then dehydrated with ethanol in a gradient concentration of 30%, 50%, 70%, 80% and 90% at room temperature for 2 h each step then incubated in 100% overnight in at 4°C. After dehydration, the samples were washed with n-Hexan for 2 h which was replaced with a clearing solution (1-part benzyl alcohol in 2-parts benzyl benzoate) without air exposure. The clearing solution was changed 2-3 times every 2 h until the samples were transparent and suitable for LSFM imaging. Samples were kept in the clearing solution at 4°C for long-term storage.

### LSFM setup, data acquisition and analysis

In the home-built LSFM setup, a customized fiber-coupled laser combiner (BFI OPTiLAS GmbH, Groebenzell, Germany) was used to provide the required excitation lines of 491, 532, 642, and 730 nm. The fluorescence was spectrally filtered by typical emission filters (AHF Analysentechnik, Tübingen, Germany) according to the use of the following fluorophores: BrightLine HC 525/50 (Autofluorescence), BrightLine HC 580/60 (A532), HQ697/58 (A647), BrightLine HC 785/62 (A750). Multicolor stacks were acquired in increments of 2 μm by imaging each plane in all color channels sequentially. Hardware components for image acquisition (laser, camera, lenses) were controlled by IQ 2.9 software (Andor, Belfast, United Kingdom). Images were saved as TIFF files and analyzed using IMARIS software v8.1.1 (Bitplane AG, CA, USA). When required, background subtraction was applied in accordance with the diameter of the cell population to eliminate unspecific background signals.

### Statistical and data analysis

Data were checked for normal or non-normal distribution. If data was normally distributed, an unpaired t-test (to compare differences between two groups) or one-way and two-way analysis of variance (ANOVA) test (to compare differences between more than two groups) were used. Non-normally distributed data were compared using the Mann-Whitney-Test. Survival between groups was compared using the log-rank (Mantel–Cox) test. All error bars indicate standard deviation. All data analyzed were from a minimum of three biological replicates (unless otherwise stated), with data from all replicates included in the statistical analysis, with no data being excluded. Statistical comparisons were made using GraphPad Prism 7 software (GraphPad). Data reaching statistical significance is indicated as: *, *p* ≤ 0.05; **, *p* ≤ 0.01; ***, *p* ≤ 0.001; ****, *p* ≤ 0.0001.

## Lead contact

Further information and requests for resources and reagents should be directed to and will be fulfilled by the Lead Contact, Andreas Beilhack

## Materials availability

This study did not generate new unique reagents.

## Data and code availability

This study did not generate new unique reagents. The data discussed in the manuscript has no restrictions on data availability. RNA sequencing data have been deposited in NCBI’s Gene Expression Omnibus and are publicly available as of the date of publication. The data are accessible through GEO Series accession number GSE291042 using the reviewer token alipqwsqflqjdsx. This study does not report original code. Any additional information required to reanalyze the data reported in this paper is available from the lead contact upon request.

Further information and requests for resources and reagents should be directed to the lead contact, Andreas Beilhack.

## Acknowledgments

This work was supported by the Deutsche Forschungsgemeinschaft (DFG; German Research Foundation, GRK2157 (Project P1, Grant No. 270563345; SFB1583 (Projects A04, A06, C03, C04, INF, Nr. 492620490); TRR124 (Projects A02, A03, A07, INF, Nr. 210879364) and TRR221 (Project Z02, Nr. 396583012). We thank Thomas Wesoly for his contribution to constructing the *S. aureus* US300 JE2 RFP-reporter strain in the laboratory of Prof. Knut Ohlsen at the Institute for Molecular Infection Biology, University of Würzburg, Germany. We are grateful to Dr. Simone Reu-Hofer (Institute of Pathology, Julius-Maximilians-University Würzburg, Germany) for her help with lung histology, to Sarah Reichardt (Helmholtz Institute for RNA-based Infection Research) for her help with RNA extraction, and to the Core Unit Systems Medicine at the University of Würzburg for excellent technical support and cDNA library preparation and sequencing. The Core Unit Systems Medicine is partly funded via project Z-6 by the Interdisciplinary Center for Clinical Research (IZKF), Würzburg. Both the scanning electron microscope JEOL JSM-7500F and the structured illumination microscope (SIM) Zeiss Elyra S.1 were funded by the DFG (Grant No. 218894895 INST 93/761-1 FUGG and No. 261184502, INST 93/823-1 FUGG, respectively). Finally, we thank Prof. Andreas Hocke and Dr. Diana Fatykhova, Charité, University medicine, Berlin, Germany) for their guidance and support with the processing of human lung samples.

## Author contributions

All authors contributed to designing the study, analyzing and interpreting data, and editing the manuscript. D.S. and A.B. wrote the manuscript. A.B. supervised the study.

