## Supplemental figures for "M-CSF–stimulated alveolar macrophages safeguard from invasive aspergillosis"

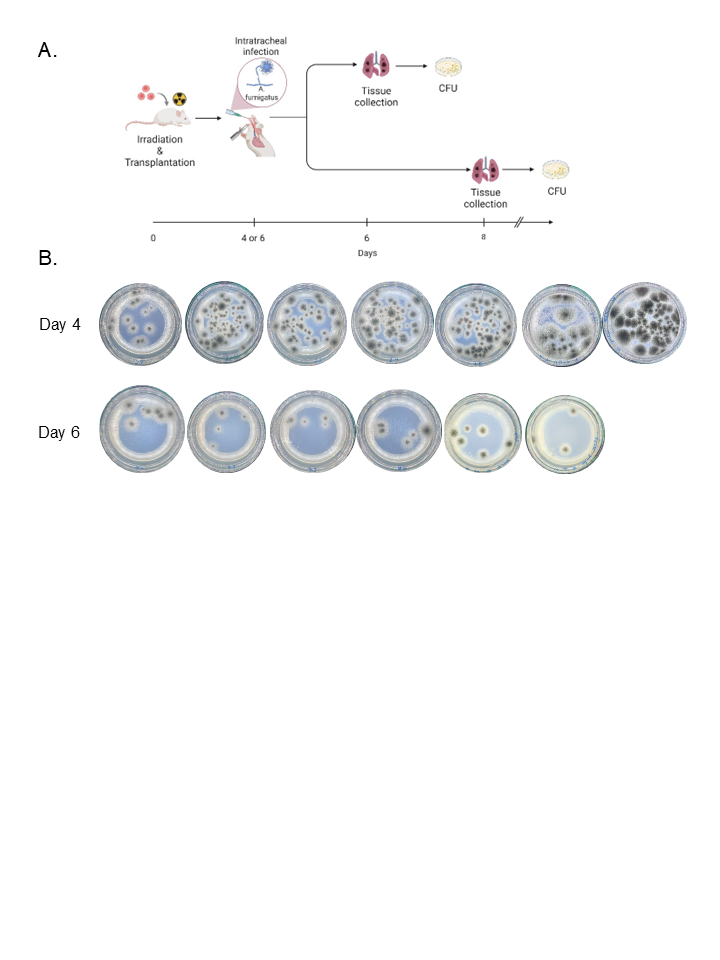

**S. figure 1: CFU assay of lung fungal burden after *A. fumigatus* infection, related to Figure 1**

**A** Experimental scheme for *A. fumigatus* intratracheal infection after irradiation with a dose of 8 Gy and allo-HCT of 5x10^6^ BM cells. The infectious dose is 5x10^4^ ATCC46645 *A. fumigatus* spores per mouse at day 4 or day 6 after allo-HCT. **B** Representative AMM agar plates showing fungal colonies recovered from lung homogenates of mice infected at day 4 or day 6 after allo-HCT. *n* = 6-7 per group.

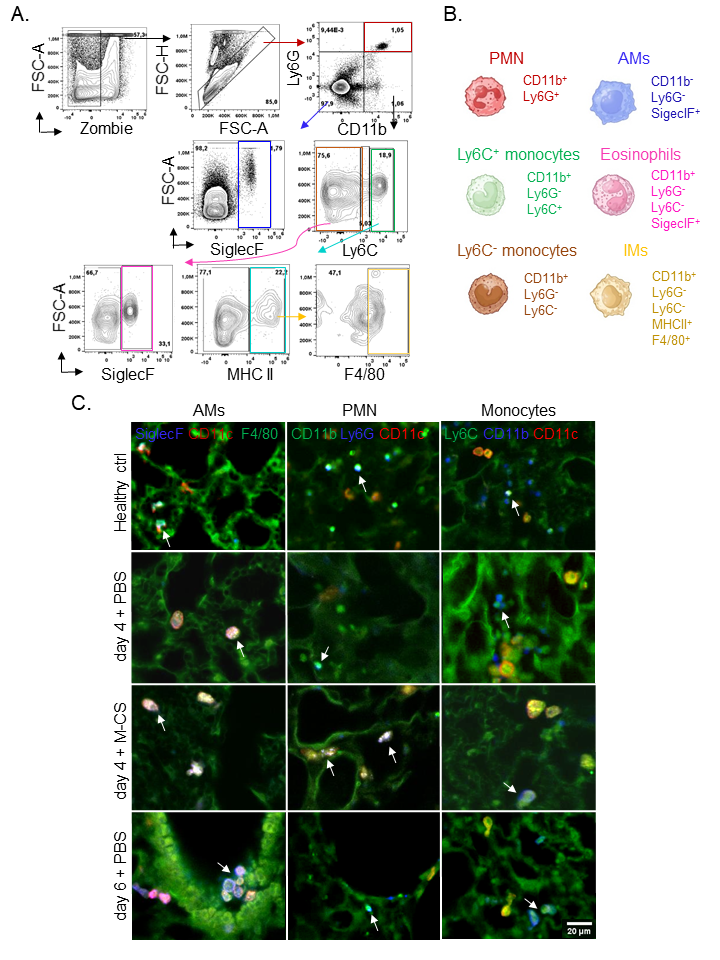

**S. figure 2: Identification of pulmonary innate immune cells in the context of *A. fumigatus* infection, related to Figure 1**

**A** Representative contour plots for the gating strategy used to identify lung innate immune cells after *A. fumigatus* infection and M-CSF treatment. **B** Representative cartoon of pulmonary innate immune cells with their differentially expressed surface markers used for their identification. **C** LSFM images of AMs, neutrophils and monocytes staining in the lungs of healthy mice and mice infected 4 and 6 days after irradiation and allo-HCT -/+ intravenous administration of M-CSF in 3 doses with a single dose of 4x10^4^ U/120 sterile H_2_O at -1, 5, 20 h of allo-HCT. Arrows indicate single stained cells. Images were taken 72 h after infection. *n* = 3 per group. Scale bar = 20 μm

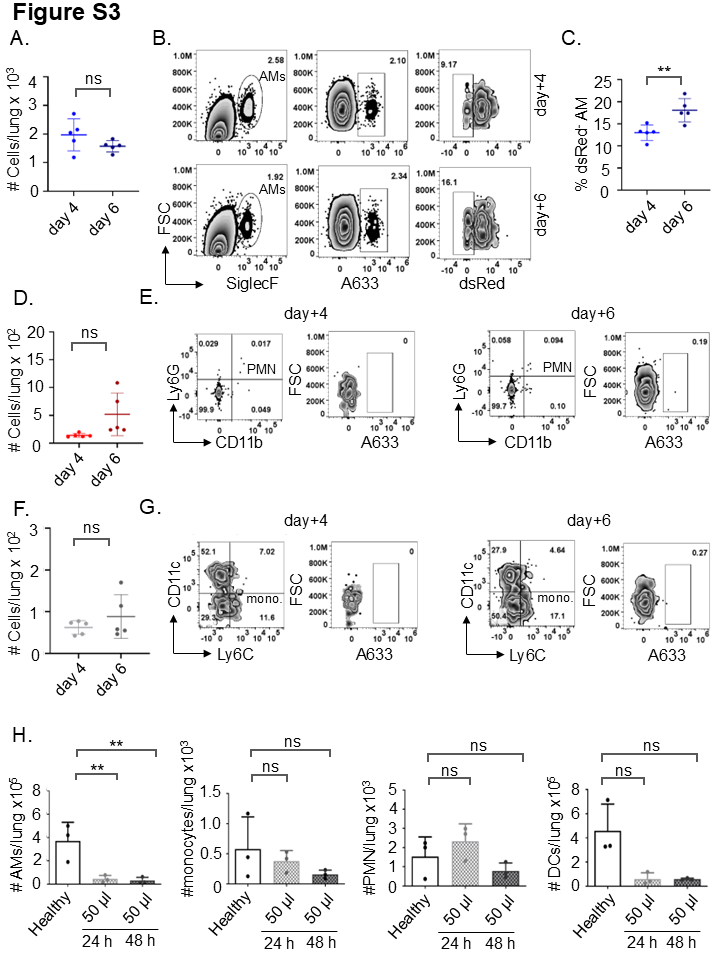

**S. figure 3: Enhanced fungal killing capacity of AMs at day 6 but not day 4 after transplantation, related to Figure 1**

**A, D, F** Flow cytometry quantification of absolute numbers of AMs, PMN, and monocytes 6 h after *A. fumigatus* infection, respectively. We infected the mice with a dose of 5x10^4^ ATCC46645 spores per mouse at day+4 and day+6 after irradiation, allo-HCT. n = 5 per group. **B** Contour plot for flow cytometry identification of AMs, which we identified by gating on single cells then living cells followed by SiglecF expression. Representative contour plot for phagocytosis and killing of FLARE conidia by AMs at 6 h after intratracheal infection with 5x10^4^ FLARE conidia/mouse both at day+4 (upper panel) and day+6 (lower panel) after transplantation based on intrinsic expression of dsRed viability signal and surface expression of Alexa Flour 633 (AF633). **C** Flow cytometry quantification of killing of FLARE conidia at 6 h after infection. We calculated the percentage of AMs involved in fungal killing by dividing the number of AMs dsRed^-^ signal per the total number of AMs wit A633+ signal. *n* = 5 per group. **E, G** Representative contour plot for phagocytosis of FLARE conidia by PMN and monocytes, respectively, at 6 h after intratracheal infection with 5x10^4^ FLARE conidia/mouse both at day+4 (left panel) and day+6 (right panel) after transplantation based on surface expression of Alexa Flour 633 (AF633). **H** Graphs for flow cytometry quantification of immune cell count in BAL fluid 24 or 48 h after intratracheal administration of 50 μl of clodronate liposomes. Values were displayed as mean ± SD. Statistical significance was determined by unpaired *t*-test (*, *p* ≤ 0.05; **, *p* ≤ 0.01). *n* = 3 per group. Values were displayed as mean ± SD. Statistical significance was determined by unpaired *t*-test (**, *p* ≤ 0.01).

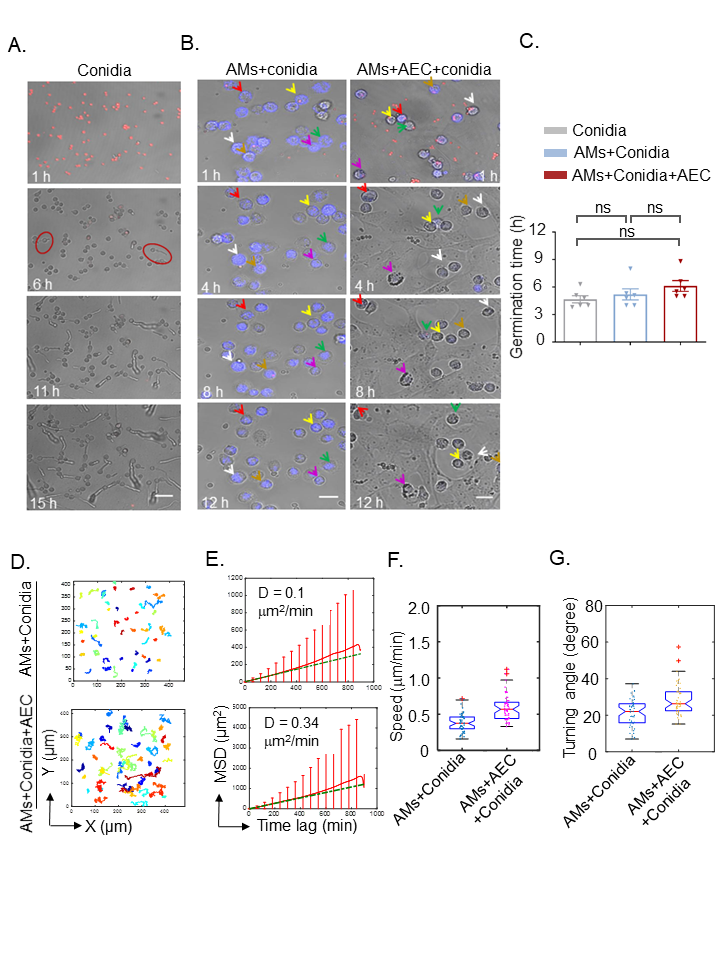

**S. figure 4: AEC prolong germination time of fungal conidia and enhance migration speed and turning angle, related to Figure 2**

**A** Confocal time-lapse imaging of the germination of *A. fumigatus* tdTomato spores over 15 h. Circles surround some germinating conidia. Scale bars = 20 μm **B** Confocal time lapse imaging for the migration of siglecF stained AMs and co-cultured with *A.fumigatus* tdTomato fungal spores with/without underlying AEC for 12 h. Scale bars = 20 μm. Colored arrows follow the migration of single AMs over time where each colored arrow indicated one AMs. **C** A graph for the germination time of tdTomato *A. fumigatus* spores with/without underlying AEC cultured together with AMs. We obtained these results by quantification of germination time in 6 different experiments. Statistical significance was determined by ordinary one-way ANOVA with Holm-Sidak’s multiple comparisons test. Values were displayed as mean ± SD. (ns, not significant). D Display of the migration path of AMs in (B). Number of cells used for analysis is 40-50 cells. **E** Quantification of MSD and diffusion coefficient of AMs based on AMs migration in (B). Number of cells used for analysis is 40-50 cells. **F, G** Quantification of the migration speed (left) and turning angle distribution (right) of AMs in (A). Number of cells used for analysis is 40-50 cells.

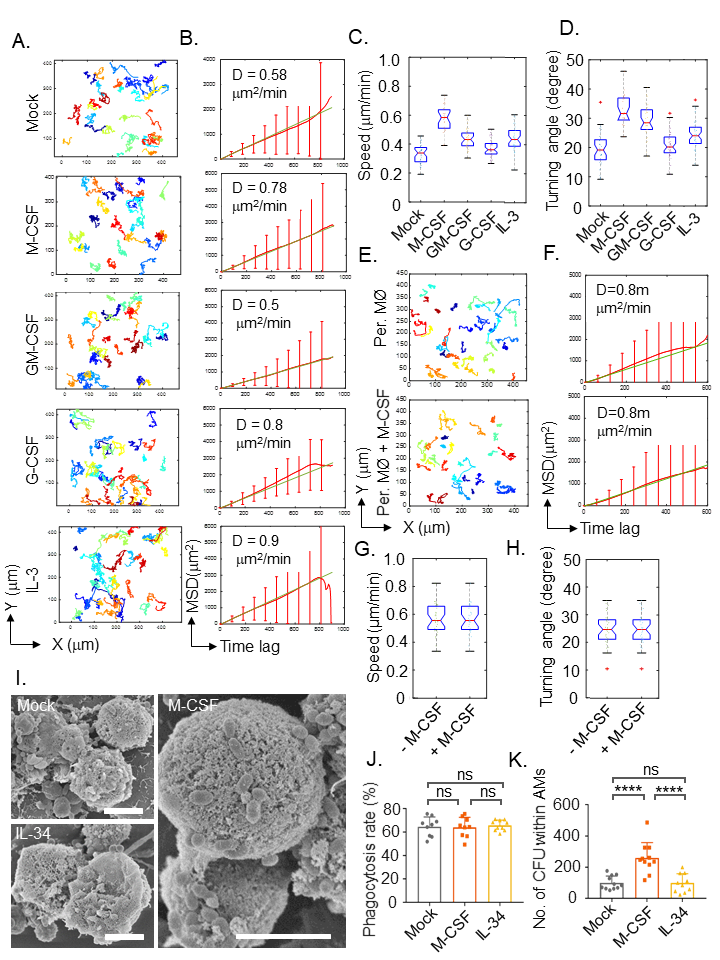

**S. figure 5: Impact of in vitro cytokine stimulation on the migration behavior and the phagocytic capacity of AMs, related to Figure 2**

**A** Display of the migration path of cytokine pre-stimulated AMs, isolated from BALB/c mice, while co-cultured with *A. fumigatus* ATCC46645 spores and underlying AEC. We performed imaging with confocal microscope for 15 h. AMs were stimulated with 300 U/mL of M-CSF, 3x10^3^ U/mL of IL-34, 6x10^3^ U/mL of GM-CSF and 6x10^3^ U/mL of G-CSF for 48 h prior to imaging. Number of cells used for analysis is 40-50 cells. **B** Quantification of MSD and diffusion coefficient of AMs based on AMs migration in (A). Number of cells used for analysis is 40-50 cells. **C, D** Quantification of the migration speed (C) and turning angle distribution (D) of AMs from (A). Number of cells used for analysis is 40-50 cells. **E** Display of the migration path of M-CSF pre-stimulated per MØ while co-cultured with *A. fumigatus* ATCC46645 spores. We performed imaging with confocal microscope for 15 h. Per MØ were stimulated with M-CSF for 48 h prior to imaging. Number of cells used for analysis is 25-40 cells. **F** Quantification of MSD and diffusion coefficient of Per MØ based on their migration in (E). Number of cells used for analysis is 25-40 cells. **G, H** Quantification of the migration speed (G) and turning angle distribution (H) of Per MØ from (E). Number of cells used for analysis is 25-40 cells. **I** Representative SEM images for AMs co-cultured with ATCC46645 conidia and AEC for 1 h in vitro with MOI of 1:5. AM are isolated by BAL and stimulated with cytokines for 48 h prior to co-culture with conidia. Scale bar = 5 µm **J, K** Graphs for flow cytometry analysis of phagocytosis rate (J) and number of CFUs (K) within AMs following overnight culture of lysed AMs on LB agar plates. (1-1.5x10^5^) AMs co-cultured with *Staphylococcus aureus* USA300_JE2 and AEC for 1 h in vitro with MOI of 1:5 followed by 1 h of extracellular killing then 1 h for intracellular killing. AMs were isolated by BAL and stimulated with cytokines for 48 h prior to co-culture with *S. aureus*. *n* =5. Values were displayed as mean ± SD. Statistical significance was determined by ordinary one-way ANOVA with Holm-Sidak’s multiple comparisons test. (*, P ≤ 0.05, **, *p* ≤ 0.01, ***, *p* ≤ 0.001).

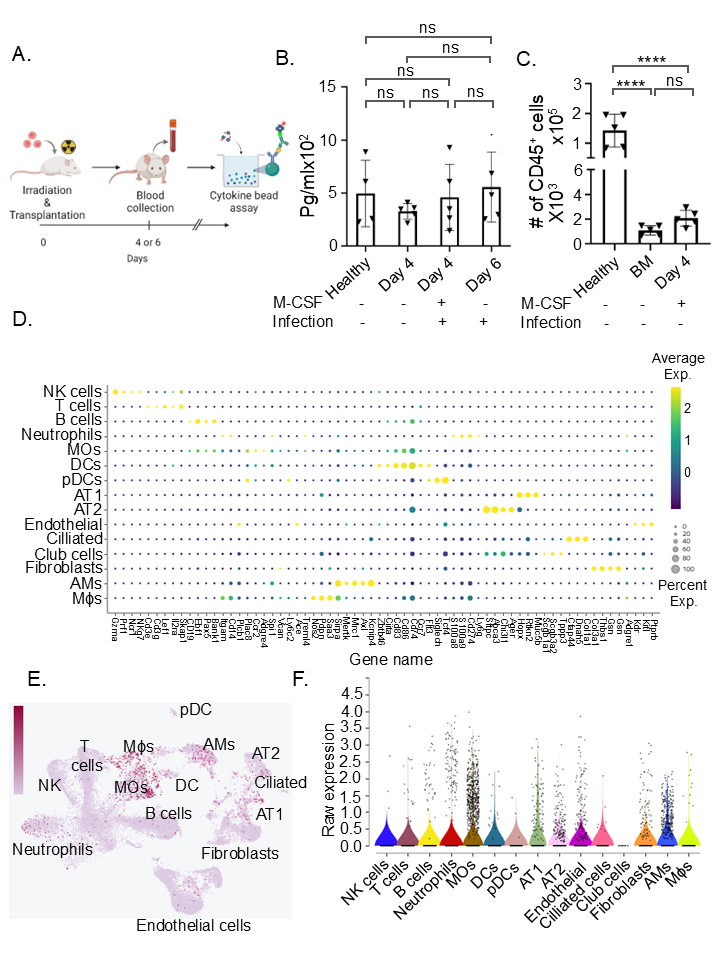

**S. figure 6: M-CSF serum levels and cellular expression of CSFR1 in the lungs, related to Figure 5**

(**A**) Experimental scheme for mice irradiation with a dose of 8 Gy and allo-HCT of 5x10^6^ BM cells. For cytokine measurement, blood was taken at day 4 or day 6 after allo-HCT. (**B**) Cytokine bead assay quantification of serum M-CSF levels at at day 4 or day 6 after allo-HCT. (**C**) Flow cytometry quantification of M-CSF effect on BM CD45^+^ cells at day 4 after allo-HCT. Mice were irradiated with a dose of 8 Gy and received allo-HCT of 5x10^6^ BM cells. M-CSF was intravenously injected in 3 doses with a single dose of 4x10^4^ U/120 µl sterile H_2_o at -1, 5, 20 h of allo-HCT *n* = 5 per group. Values were displayed as mean ± SD. Statistical significance was determined by ordinary one-way ANOVA with Holm-Sidak’s multiple comparisons test. (ns, non-significant, *, P ≤ 0.05, **, *p* ≤ 0.01, ***, *p* ≤ 0.001, ****, *p* ≤ 0.0001). (**D**) Single-cell RNA sequencing of lung cells from uninfected mice was carried out using the Evercode cell fixation kit and WT kit v3 and data were analyzed with Trailmaker (Parse Biosciences). The dot plot shows marker gene expression used for cell-type annotation. The colour gradient represents the mean expression level of each gene within a cell type, and the dot size indicates the percentage of cells expressing that marker. AM: alveolar macrophages; MOs:monocytes; Mϕs:macrophages; DC: dendritic cells; pDC: plasmacytoid dendritic cells; AT1/2: alveolar type 1/2 cells; ciliated: ciliated epithelial cells; NK: natural killer cells. (**E**) Uniform Manifold Approximation and Projection (UMAP) plot displaying the annotated cell types, with the purple colour gradient indicating the expression level of *Csf1r*. (**F)** Violin plot showing the expression of *Csf1r* transcripts in individual cells of each type. Values represent normalized raw transcript counts per cell, as visualized by the default raw expression module

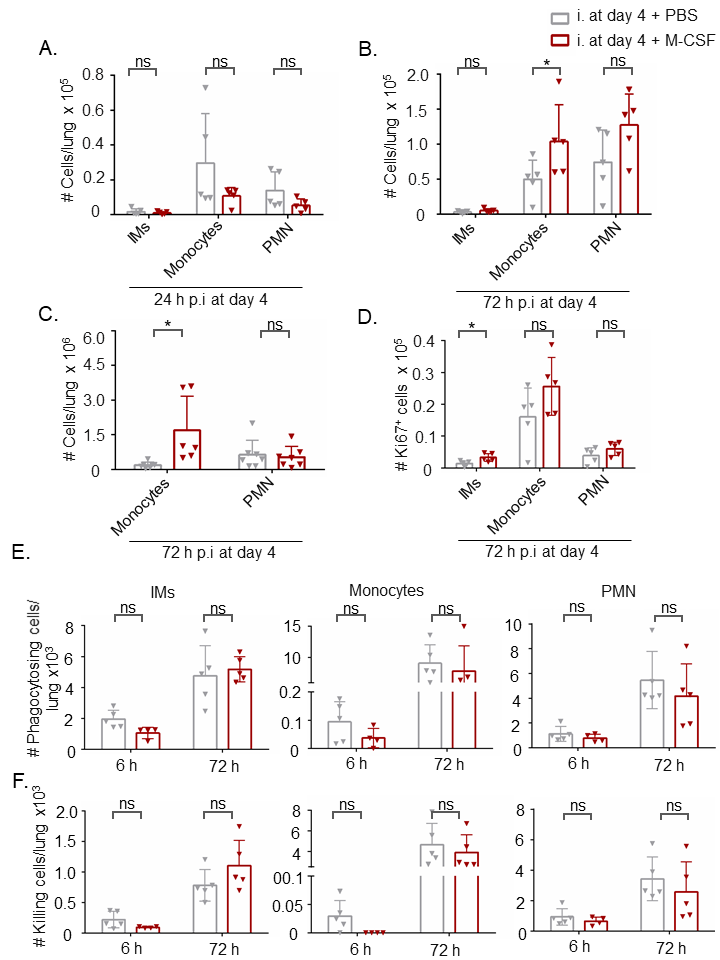

**S. figure 7: The effect of exogenous M-CSF on proliferation and phagocytic function of different local pulmonary immune cell subsets, related to Figure 5**

**A, B** Flow cytometry quantification of IMs, neutrophils and monocytes numbers in the lungs at 24 h (A) and 72 h (B) after *A. fumigatus* infection. We infected the mice with a dose of 5x10^4^ ATCC46645 spores per mouse at day 4 after irradiation, allo-HCT and -/+ M-CSF in vivo administration. n = 5 per group. **C** A graph for LSFM quantification of monocytes and neutrophils number at 72 h after *A. fumigatus* infection. We quantified 3 different z stacks, each with a size of 250-400 µm^3^, per each mouse with total of 3 mice per condition. **D** Flow cytometry analysis of IMs, neutrophils and monocytes proliferation via Ki67^+^ intracellular staining at 72 h after *A. fumigatus* infection. *n* = 5 per group. **E, F** Flow cytometry quantification of absolute numbers of AMs involved in fungal phagocytosis (E) and fungal killing (F) at 6 h and 72 h after *A. fumigatus* infection. *n* = 5 per group. Values were displayed as mean ± SD. Statistical significance was determined by unpaired *t*-test (*, *p* ≤ 0.05).

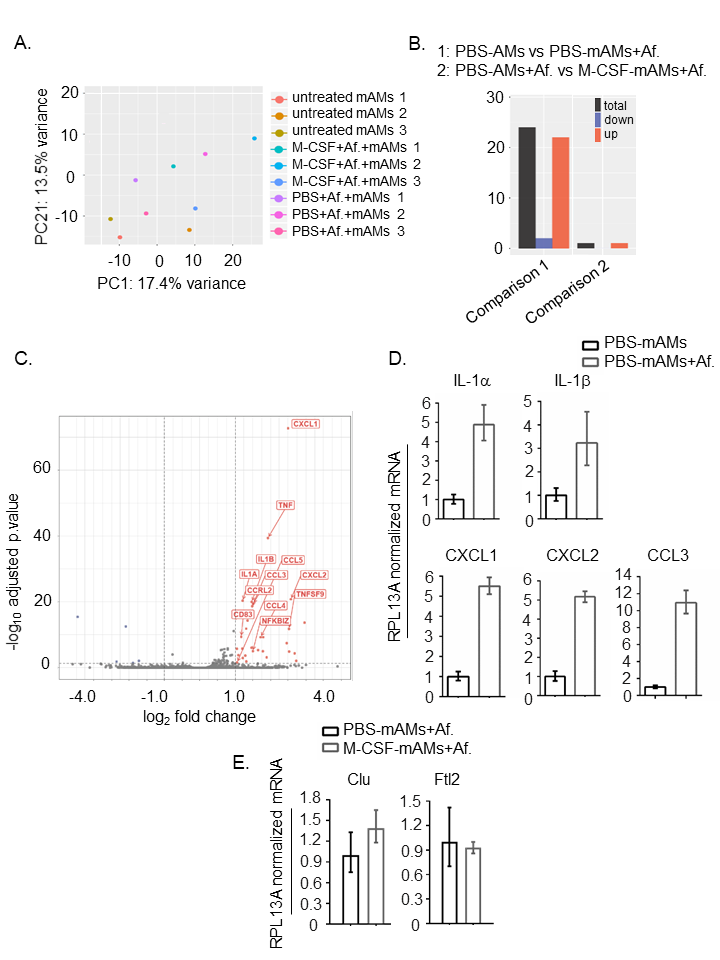

**S. figure 8: RNA-seq of differentially stimulated AMs co-cultured with ATCC46645 spores, related to Figure 5**

**A** PCA of AMs after co-culture for 6 h in vitro with *A. fumigatus* ATCC46645 spores in MOI 1:5. The analysis is based on 3 different biological replicates per group. *n* = 3 per group. **B** Representative bar graph for the total numbers of DEGs of PBS-stimulated AMs. RNA was isolated from AMs after in vitro co-culture with ATCC46645 spores for 6 h with MOI 1:5. AMs were isolated by BAL and stimulated with cytokines for 48 h prior to co-culture with fungal spores. The analysis is based on 3 different biological replicates per group. n = 3 per group. **C** A volcano plot represents DEGs resulted from the comparison between PBS-stimulated AMs vs. PBS-stimulated AMs co-cultured with ATCC46645 spores for 6 h with MOI 1:5. n = 3 per group. **D** Representative bar graphs for qPCR confirmation of DEGs by PBS–stimulate AMs. We obtained cDNA after RNA isolation from AMs, which were co-cultured in vitro for 6 h with MOI 1:5 of *A. fumigatus* ATCC46645 spores. mRNA level was normalized to the level of the house keeping gene RPL13A. AMs were isolated by BAL and stimulated them with cytokines for 48 h prior to co-culture with the fungus. Quantification is based on comparative threshold cycle method (2^-∆∆CT^ method) using 3 different biological replicates per group where for each biological replicate 3 technical replicate. **E** Representative bar graphs for qPCR confirmation of DEGs by M-CSF–stimulate AMs. We obtained cDNA after RNA isolation from AMs, which were co-cultured in vitro for 6 h with MOI 1:5 of *A. fumigatus* ATCC46645 spores. mRNA level was normalized to the level of the house keeping gene RPL13A. AMs were isolated by BAL and stimulated them with cytokines for 48 h prior to co-culture with the fungus. Quantification is based on comparative threshold cycle method (2^-∆∆CT^ method) using 3 different biological replicates per group where for each biological replicate 3 technical replicate.
